# Reduced myofibroblast transdifferentiation and fibrotic scarring in ischemic stroke after imatinib treatment

**DOI:** 10.1101/2021.10.28.466225

**Authors:** M. Zeitelhofer, C. Stefanitsch, J. Protzmann, D. Torrente, M.Z. Adzemovic, S.A. Lewandowski, L. Muhl, U. Eriksson, I. Nilsson, EJ. Su, D. A. Lawrence, L. Fredriksson

## Abstract

The tyrosine kinase inhibitor imatinib has been reported to improve outcome in patients following ischemic stroke but the exact mechanism remains elusive. Here, utilizing a photothrombotic murine model of middle cerebral artery occlusion (MCAO), we show that imatinib-mediated inhibition of stroke-induced blood-brain barrier (BBB) dysfunction coincided with decreased expression of genes associated with inflammation and fibrosis in the cerebrovasculature. We found that imatinib effectively dampened stroke-induced reactive gliosis and myofibroblast transdifferentiation, whilst having very limited effect on the rest of the glia scar and on peripheral leukocyte infiltration. Further, our data suggest that consolidation of the PDGFR*α*^+^ portion of the fibrotic scar in imatinib-treated mice contributes to the improvement in functional outcome compared to the vehicle controls, where the PDGFR*α*^+^ scar is expanding. Comparison with human stroke transcriptome databases revealed significant overlap with imatinib-regulated genes, suggesting imatinib may modulate reactive gliosis and fibrotic scarring also in human stroke.

## Introduction

Ischemic stroke represents a major public health challenge and is currently the 3rd leading cause of death and the leading cause of adult neurological disability in the Western world^1^. Further, stroke has been reported in 2–6% of patients hospitalized with COVID-19^2^. Treatment options are limited and currently the only approved interventions are intravenous thrombolysis (IVT) with recombinant tissue plasminogen activator (tPA) and mechanical thrombectomy (MT)^3^. However, despite combined IVT and MT, up to 40% of the patients will die or remain functionally dependent, and more importantly, due to various contraindications, only a fraction of all ischemic stroke patients will be treated with these therapies. Much effort has therefore been invested in researching additional treatment options, mainly focusing on direct neuroprotection. However, due to the limited success of neuroprotection approaches, studies examining the therapeutic potential of preserving or restoring the integrity of the blood-brain barrier (BBB) have been gaining interest^4^. This is supported by our published study showing for the first time that a treatment strategy aiming to reduce BBB damage using the tyrosine kinase inhibitor imatinib improves neurological outcome in humans suffering from ischemic stroke^5^. Despite these advances the dynamics and functional relationship of cerebrovascular dysfunction with ischemic stroke progression remain largely unclear and require further investigation.

The BBB forms a mechanical and functional barrier between the systemic circulation and the central nervous system (CNS) and it tightly controls trafficking of substances between the blood and the CNS^4^. The barrier properties are established by the brain endothelial cells. It is however widely recognized that other cells in the neurovascular unit (NVU), including perivascular astrocytes and vascular mural cells (vascular smooth muscle cells (vSMC) and pericytes) as well as neurons and microglia, all work together in a coordinated way to maintain BBB properties^4^. Damage to the BBB is an early pathological event in ischemic stroke, and the absence of a functional BBB will lead to profound disturbances in neuronal and glial signaling^6^. Several molecular pathways have been reported to play important roles in disease-induced BBB damage, including our previous studies in mice demonstrating the role of tPA-activated platelet-derived growth factor CC (PDGF-CC) signaling via its tyrosine kinase receptor PDGF receptor α (PDGFRα) on perivascular cells in the NVU^7–13^. As aforementioned, administration of imatinib, a small molecule tyrosine kinase inhibitor of Abl, c-Kit and PDGFR^14^, significantly improves outcome after both ischemic and hemorrhagic stroke in rodents^7, 15, 16^ and in humans^5^. This beneficial effect has been ascribed to imatinib’s potential to reduce stroke-induced BBB leakage, but how this exactly translates to reduced lesion volume and improved neurological outcome is yet unknown. However, activation of the endothelium and subsequent BBB damage in the acute phase (hours after onset) of cerebral ischemia is known to accelerate secondary degeneration of the surrounding, initially spared, neural tissue, and this continues during the subacute phase (hours – days after onset)^17^. In addition, cells in the NVU act as sensors for brain injury eliciting activation of the early reactive gliosis response, in which astrocytes, NG2 glia (also referred to as oligodendrocyte progenitor cells (OPCs) or polydendrocytes) and microglia become activated, leukocytes infiltrate and a whole cascade of post stroke neuropathology is initiated^18^.

The general view is that reactive gliosis is an important early injury response, orchestrating subsequent formation of a glial scar during the tissue remodeling phase after insult (days – weeks after onset)^19^. It is believed that demarcation of the lesion by activated glia cells sequesters the toxic environment at the lesion site, thereby protecting the relatively unaffected surrounding CNS tissue. Also, isolation of the lesion is thought to allow regeneration of the injured tissue and thus enable recovery of CNS function in the chronic phase after insult. However, the role of the scar is highly debated and sustained gliosis has been reported to be deleterious to functional recovery^20^. For example, reactive gliosis and glial scarring, especially astrogliosis, are regarded as barriers to CNS regeneration and have been shown to have inhibitory effects on CNS axon regrowth^21^. Thus, these discrepancies highlight the need for a deeper understanding of the CNS injury response and repair.

It has been proposed that CNS responses to injury resemble that of normal wound healing processes in other tissues and that the poor CNS recovery is reminiscent of chronic/unresolved wounds^17^. A key healing phase in normal wound healing is ‘scar contraction’, which is achieved by injury-induced transdifferentiation and proliferation of specialized subsets of mesenchymal cells, mainly localized in a perivascular position, into contractile myofibroblasts^22^. These myofibroblasts contribute to repair by generating contractile forces in the tissue and are identified by expression of *α*-smooth muscle actin (ASMA), a marker not normally expressed in tissue mesenchymal cells at steady state. It is not yet clear if a similar process occurs in CNS lesions; however, hypoxia has been shown to promote myofibroblast transition in other organs^23^. Although beneficial initially, persistent myofibroblast expansion triggers pathological tissue contraction and results in overproduction of extracellular matrix (ECM), a pathology known as fibrosis^22^. This contributes to distortion of parenchymal architecture, which compromises organ recovery and impairs function. Interestingly, PDGF-CC/PDGFRα signaling has been associated with the fibrotic response in other organs^24–28^ and nintedanib, a tyrosine kinase inhibitor that blocks the activity of PDGF receptors as well as that of FGF and VEGF receptors, has been shown to inhibit myofibroblast transdifferentiation and proliferation in patients with idiopathic pulmonary fibrosis^23^.

In the present study, we utilized a photothrombotic murine model of experimentally induced ischemic stroke in order to elucidate the molecular and cellular mechanisms of imatinib treatment. We found that stroke-induced BBB leakage was inhibited by imatinib and coincided with preserved cellular organization in the NVU in the acute phase after ischemia onset. BBB transcriptome analyses of vascular fragments isolated from the ipsilateral hemisphere of imatinib-treated mice and their controls identified differential gene expression mainly associated with inflammation and fibrosis pathways. Immunohistological analyses confirmed that imatinib dampened the reactive gliosis response and myofibroblast transdifferentiation after stroke, whilst having very limited effect on peripheral leukocyte infiltration and on the rest of the glial scar. Assessment of sensory-motor integration after stroke revealed that functional benefit with imatinib treatment progressively improved over time. This suggests that imatinib might ameliorate stroke outcome by counteracting the spread of fibrosis without affecting the ability of the rest of the glial scar to protect the healthy parenchyma from the toxic environment of the lesion. This study also offers novel insight into the relationship between BBB breach and fibrotic/glia scarring as well as the role of myofibroblasts in CNS injury response and repair.

## Results

### Imatinib attenuates cerebrovascular leakage and preserves normal organization of the NVU in the acute phase after MCAO

We have previously shown that tPA-mediated activation of PDGF-CC/ PDGFR*α* signaling in the NVU during ischemic stroke in mice induces opening of the BBB and augments brain injury^7^. Blocking this pathway with the tyrosine kinase inhibitor imatinib improves outcome following ischemic stroke in both mice^7^ and human patients^5^ as well as in other neuropathologies in mice^8–12, 15^. However, very little is known about the kinetics and the underlying mechanisms of this inhibition. To advance this knowledge we performed *in vivo* BBB leakage, BBB transcriptomics and immunofluorescent analyses at different time points following MCAO induction in mice treated with imatinib or PBS as vehicle control (Fig. 1a, study design).

**Figure 1.**
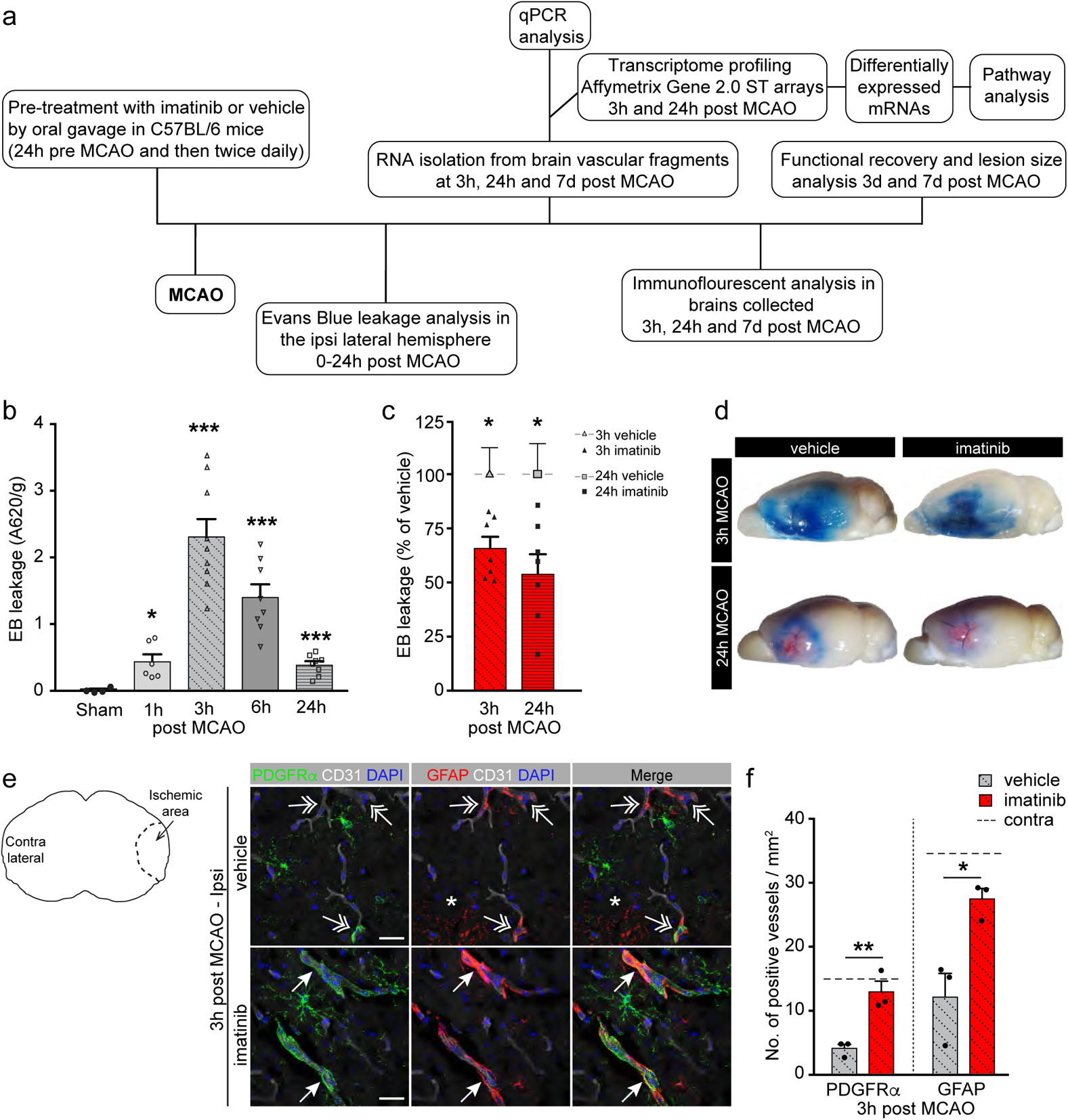
Imatinib attenuates MCAO-induced cerebrovascular breach. (a) Schematic illustration of study design. (b) MCAO was induced in vehicle-treated wild type mice and Evans blue (EB) dye was injected intravenously 1 hour before transcardial perfusion with PBS. Cerebrovascular permeability was quantified by EB extravasation analysis in the ipsilateral ischemic hemisphere at various time points post MCAO. Increased EB extravasation was detected already 1 hour post MCAO and peaked at 3 hours post MCAO. Significance as compared to sham operated controls. (c) Cerebrovascular permeability in the ipsilateral ischemic hemisphere of imatinib-treated mice was significantly decreased at 3 and 24 hours post MCAO compared to the respective vehicle controls (normalized to 100%). (d) Representative images of EB extravasation in brains from vehicle and imatinib-treated mice 3 and 24 hours post MCAO. (e) Immunofluorescent co-staining for PDGFR*α* (green), GFAP (red), and the endothelial cell marker CD31 (white) was performed on brain sections from imatinib and vehicle-treated mice 3 hours post MCAO. Imatinib treatment preserved high expression of PDGFR*α* and GFAP in perivascular cells around vessels (arrows) in the ischemic area (see schematic illustration) whereas it was lost or scattered (two headed arrows) in vehicle controls. Non-vascular GFAP signal (asterisk) was however increased in vehicle controls. Representative maximum intensity projections of 8 µm confocal z-stacks are shown. Cell nuclei were visualized by DAPI (blue). Scale bar, 25 µm. (f) Quantification of PDGFR*α* and GFAP positive vessels based on counting positive vessels in eight confocal images per brain in the ischemic area. Dashed lines represent contralateral group mean. Symbols represent individual data points and bars the group mean ± S.E.M. Statistical significance *p < 0.05; **p < 0.01; ***p < 0.001 relative to control (b, one-way ANOVA with Welch’s correction; c, f, Student’s unpaired t-test).

To study the kinetics of imatinib attenuated BBB permeability, we first determined the time course of MCAO-induced BBB leakage during the first 24 hours after ischemic onset in our photothrombotic model (Fig. 1b). Evans blue (EB) extravasation analyses revealed a significant, time-dependent increase in EB extravasation in the ipsilateral hemisphere of vehicle control mice after MCAO compared to sham operated controls. We found that already 1 hour post MCAO the BBB is losing its integrity. At 3 hours post MCAO we detected the highest leakage, which was followed by a time-dependent decrease in EB extravasation, displaying a similar level of EB extravasation at 24 hours as at 1 hour post MCAO. These findings correlate well with previous reports demonstrating an early transient rise in BBB permeability in various reperfusion models of MCAO^29, 30^. To determine the kinetics of using imatinib to block MCAO-induced BBB leakage, we compared EB extravasation 3 and 24 hours post MCAO in imatinib-treated mice to vehicle controls. The results showed a significant reduction in EB extravasation of approximately 34% 3 hours and 46% 24 hours post MCAO in imatinib-treated mice compared to vehicle controls (Fig. 1c). The latter correlates well to previously published data^7^. Representative images illustrate the transient nature of MCAO-induced BBB leakage, with extensive EB extravasation 3 hours post MCAO and less at 24 hours, which is strongly attenuated by imatinib treatment (Fig. 1d).

Immunofluorescent stainings revealed that the extensive BBB breach observed 3 hours post MCAO coincided with changes in the NVU of PDGFR*α*^+^ vessels in the ischemic region of vehicle control mice, whereas imatinib treatment preserved a more normal appearing organization similar to that seen in naïve unchallenged brains (compare Fig. 1e to Extended data Fig. 1). In naïve brains, as well as in the non-ischemic contralateral hemisphere, PDGFR*α* was found in perivascular cells staining positive for aquaporin 4 (AQP4) and glial-fibrillary acidic protein (GFAP) but not in AQP4^+^ cells distributed along capillaries (Extended data Fig. 1a – c). PDGFR*α* was also detected in non-vascular NG2-glia cells (Extended data Fig. 1a, d). Further, the PDGFR*α*^+^ perivascular cells were located on the parenchymal side of the vascular smooth muscle cell (vSMC) layer in alpha smooth muscle actin (ASMA) positive vessels (Extended data Fig. 1c). In our analyses of stroked brains 3 hours post MCAO we found that imatinib treatment preserved high expression of PDGFR*α* and GFAP around vessels within the ischemic area, whereas it was absent or scattered around vessels in the ischemic area of vehicle controls (Fig. 1e). Meanwhile we noticed an increase of non-vascular GFAP signal in vehicle controls (asterisk, Fig. 1e). Quantification of PDGFR*α*^+^ and GFAP^+^ vessels in the ischemic area confirmed these observations (Fig. 1f).

This is in line with our previous published data showing that increased BBB opening after MCAO is mediated via early activation of PDGFR*α* signaling in perivascular astrocytes^7, 8, 31^ and suggests that strategies targeting BBB integrity might be most effective if provided in the early phase after ischemic stroke.

### Imatinib treatment regulates expression of genes associated with fibrosis and inflammation in the cerebrovasculature after MCAO

To further investigate imatinib’s effect after ischemia we performed transcriptome analysis on cerebrovascular fragments isolated 3 and 24 hours post MCAO from imatinib-treated mice or vehicle controls. Differential gene expression analysis was performed by microarray hybridization (GeneChip Gene 2.0 ST Array) or by qPCR. Using a *P-*value < 0.05 and a log2 (0.5) fold change as cutoff, we identified 121 and 85 differentially expressed transcripts in the cerebrovascular fragments from imatinib-treated mice at 3 hours (Supplementary table 1) and 24 hours (Supplementary table 2) post MCAO, respectively, compared to vehicle controls (Fig. 2a). Raw data are deposited on the NCBI Gene Expression Omnibus database (accession no. GSE137534). Pathway analysis with Ingenuity revealed that functions related to the immune system, vascular damage and leukocyte adhesion, as well as fibrosis were modulated by imatinib treatment compared to vehicle controls both at 3 hours (Fig. 2b) and 24 hours (Fig. 2c) post MCAO. Subsequent manual analysis of each dataset and comparison to the harmonizome databank^32^ confirmed these findings and revealed that approximately 15% of the imatinib regulated transcripts were associated with inflammation and 30% with fibrosis, including *Ccl2, Cdh11, Cntfr, Cxcl10, Hla-A, Pdgfra* and *Tmsb4x* at 3 hours post MCAO (Fig. 2d, Supplementary table 1) and *Ccl5, Ccl22, Hpse, Il1r2, Itgax* and *Pdgfd* at 24 hours post MCAO (Fig. 2f, Supplementary table 2). Most of the differentially expressed inflammation and fibrosis associated genes regulated by imatinib in the cerebrovascular fragments at 3 hours post MCAO (Supplementary table 3) were found to be expressed by perivascular astrocytes (AC) and fibroblast like cells (FB) of the unchallenged NVU according to the single cell RNAseq database of mouse brain vascular and vessel-associated cell types^33^. This supports potential involvement of these cells in inflammatory and fibrotic processes following ischemia. Comparison of our microarray data with the publicly available microarray dataset from the perihematomal area of human stroke patients (GSE24265)^34^, an area associated with high levels of edema formation, revealed high overlap and suggest that our murine analyses may also be relevant for human disease. Approximately a 45% (35 overlapping genes out of 78 genes expressed in both datasets) and a 42% (21 out of 50 genes expressed in both datasets) overlap was detected in comparison with our 3 and 24 hours datasets, respectively (Fig. 2g and h, Supplementary table 4 and 5). Considering that the human data was generated from whole tissue samples from hemorrhagic stroke patients, and our data are from cerebrovascular fragments after MCAO, this overlap is high. Although, it should be noted that imatinib has shown benefit also in experimental models of hemorrhagic stroke^15^. Interestingly, approximately half of the overlapping genes were associated with fibrosis (Fig. 2g, h and Supplementary table 4, 5).

**Figure 2.**
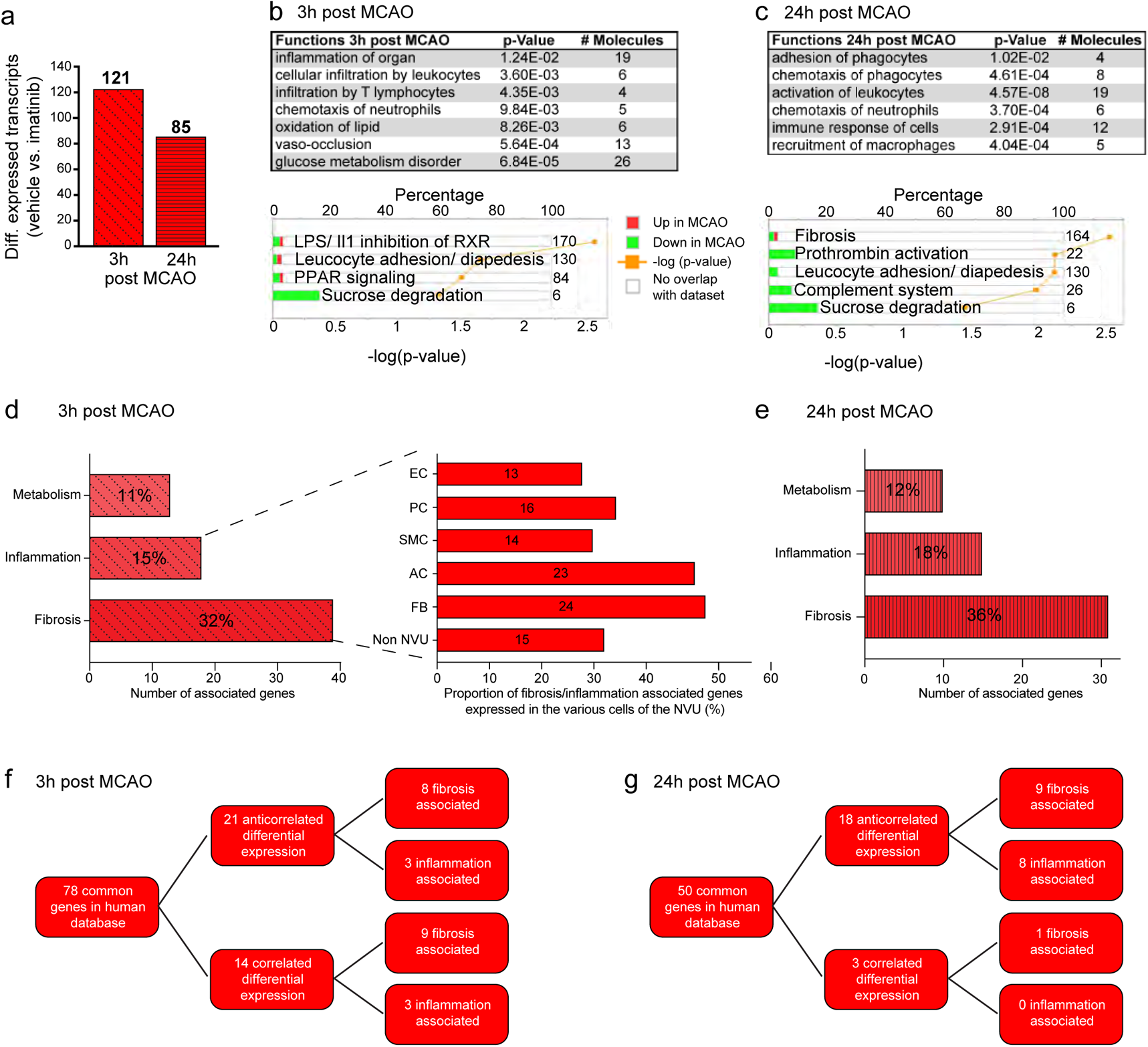
Kinetic analysis and functional clustering of the cerebrovascular transcriptome after MCAO reveals that imatinib affects expression of fibrotic and inflammatory associated transcripts. Brains from imatinib and vehicle-treated mice were harvested at various time points after MCAO or from sham operated controls (n = 2 – 3 mice / time point) and RNA was isolated from cerebrovascular fragments in the ipsilateral hemisphere using a magnetic sorting method. Gene expression analysis of the BBB transcriptome was performed using Affymetrix Mouse Gene 2.0 ST arrays. (a) Microarray analysis of vascular fragments isolated from imatinib-treated mice compared to vehicle controls revealed that 121 and 85 transcripts were differentially expressed at 3 and 24 hours post MCAO, respectively. We used a *P*-value < 0.05 and a log2 (0.5) fold change as cutoff for differentially expressed genes. (b – c) Ingenuity pathway analysis demonstrated cerebrovascular regulation of disease-relevant pathways 3 hours (b) and 24 hours (c) post MCAO in imatinib-treated mice compared to vehicle controls. The numbers to the right of the panel represent the number of molecules associated with the respective pathway. For pathway analysis, significance was determined with the right-tailed Fisher’s exact test depicted by –log(*P*-value). (d, e) Manual functional annotation and comparison with the harmonizome database showed a high percentage of the differentially expressed genes to be associated with fibrosis followed by inflammation and metabolism 3 hours (d) and 24 hours post MCAO (e). Comparison with the mouse brain single cell vascular atlas suggests that a high percentage of the differentially expressed fibrosis and inflammation associated genes at 3 hours are normally expressed in the NVU, especially in perivascular astrocytes (AC) and fibroblast like cells (FB). (f, g) Comparison of our datasets with a microarray dataset from perihematomal areas of stroke patients reveals high overlap especially with fibrosis and inflammation associated genes.

### Imatinib attenuates the reactive gliosis response within hours after MCAO

Based on our BBB transcriptomics analysis we next investigated the effect of imatinib on reactive gliosis, the initial events of fibrosis and inflammation in response to insult. Among the glial cells taking part in the reactive gliosis response are astrocytes, NG2-glia cells and microglia/macrophages, which are recruited to the site of insult^19^. Since astrocytes are known to react to injury by hypertrophy and up-regulation of GFAP in the acute phase following insult^35^, we assessed astrogliosis by immunofluorescent staining for GFAP 3 hours post MCAO. These data demonstrated that there was significantly increased GFAP signal in the ischemic area (demarcated with a dashed line) compared to the non-ischemic surrounding tissue (Fig. 3a, b). Our stainings revealed that imatinib inhibited this MCAO-induced activation of astrocytes, displaying approximately 60% less GFAP positive signal in the ischemic area compared to vehicle controls 3 hours post MCAO (quantified in Fig. 3c). This pronounced effect seen on astrocytes is even more remarkable given that imatinib treatment preserves perivascular GFAP expression around vessels (see Fig. 1e, f and arrows, Fig. 3b) and is supported by our gene profiling analyses, showing that many of the imatinib regulated genes in the cerebrovascular fragments are normally expressed in perivascular astrocytes within the unchallenged NVU (Fig. 2d, Supplementary table 3). Further analysis of the reactive gliosis response showed that NG2-glia cell condensation, determined by staining for PDGFR*α* and neuron glia antigen-2/CSPG4 (NG2), appeared in the ischemic border of vehicle control brains 3 hours post MCAO (two headed arrows) but was inhibited by imatinib treatment (Fig. 3d – f). Likewise, imatinib reduced microgliosis/macrophage recruitment after MCAO, as determined with staining for CD11b (microglia, arrows; activated microglia/macrophages, two headed arrows, Fig. 3g – h). Quantification revealed no detectable (ND) activated microglia/infiltrating macrophages 3 hours post MCAO whereas at 24 hours and 7 days post MCAO, pronounced microglia activation/macrophage infiltration was detected and this was significantly reduced by imatinib treatment (Fig. 3h). Considering our earlier findings with the CX3CR1^GFP^/CCR2^RFP^ reporter mice, showing very few infiltrating macrophages up to 24 hours post MCAO^31^, the effect of imatinib, at least up to 24 hours after MCAO, appears to be mainly mediated by inhibiting activation of intrinsic microglia. This is further supported by our gene expression analyses in Fig. 2, showing that imatinib is inhibiting proinflammatory mediators known to activate microglia. For instance, our analyses (Supplementary table 1) demonstrated that imatinib treatment led to decreased expression of *Sirpb1* and *Cntfr*, factors reported to enhance the phagocytic activity and activation of microglia^36^, whereas levels of *Cd55* and *Rnf113a1*, factors expressed by microglia that inhibit complement activation and terminate CXCR4 signaling^37^, were increased. It is of course possible that imatinib affects recruitment of macrophages in later phases after MCAO. This is supported by imatinib-induced upregulation of *Nov* expression, a factor reported to reduce monocyte adhesion. Apart from this potential effect on macrophage recruitment, our immunohistochemical analyses suggest that imatinib has only limited effect on peripheral immune cell infiltration after MCAO, including neutrophils and T cells (Extended data Fig. 2). Taken together this indicates that imatinib attenuates the reactive gliosis response after MCAO. This is supported by qPCR analyses showing that MCAO-increased expression of *Il1a, Tnfa* and *Ccl2*, known drivers of endothelial cell activation, microglia activation/macrophage infiltration as well as fibrosis, was dampened by imatinib treatment (Fig. 3i).

**Figure 3.**
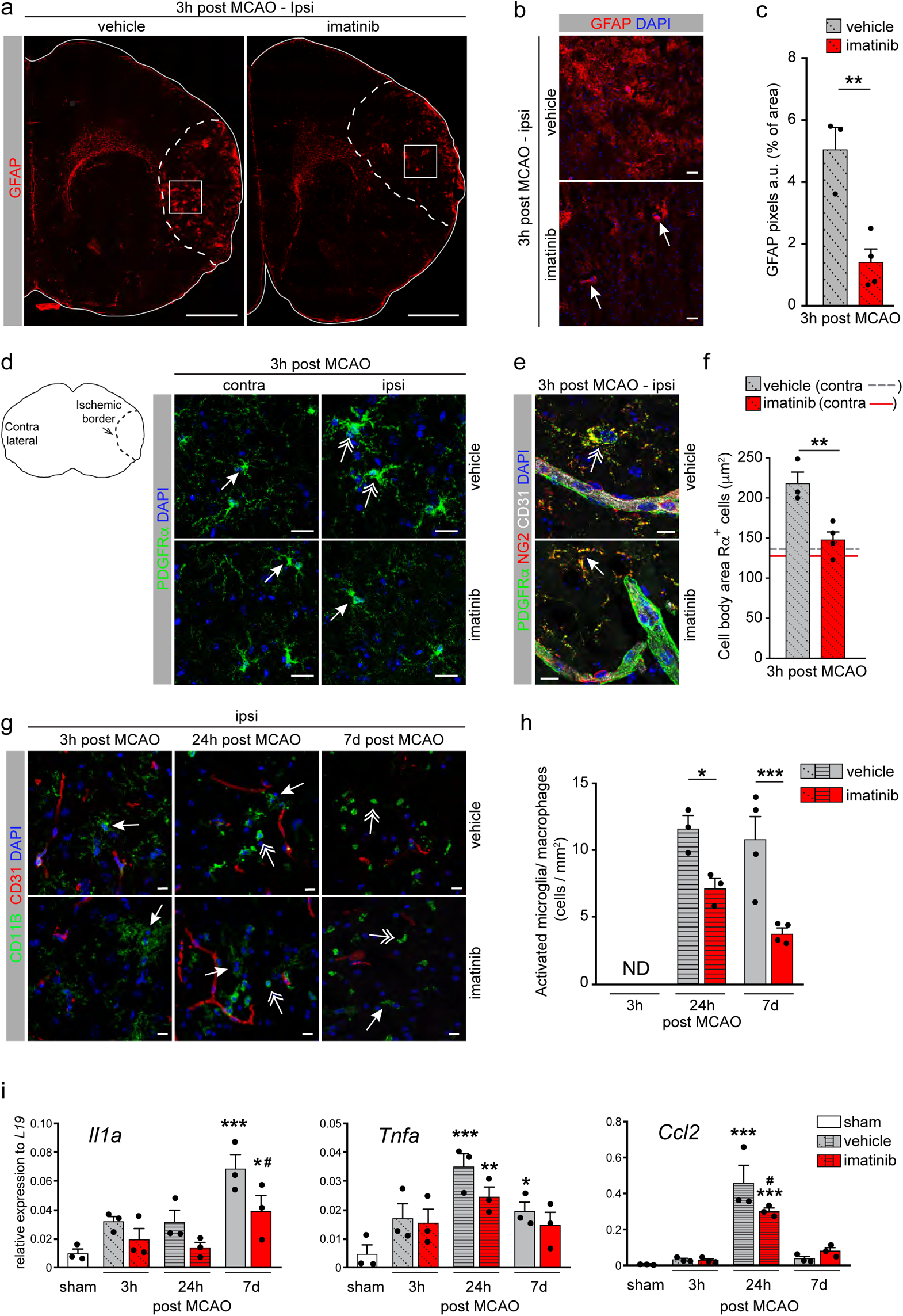
Imatinib attenuates the reactive gliosis response in the acute phase after MCAO. (a, b) Immunofluorescent staining for the astrocyte marker GFAP (red) was performed on brain sections from imatinib and vehicle-treated mice 3 hours post MCAO. (a) Stitched tiles of epifluorescent images (taken with a 10x objective). (b) Confocal images of the boxed regions in (a) show that imatinib reduced astrocyte activation in the ischemic area while preserving vascular associated GFAP^+^ staining (arrows). (c) Quantification of GFAP signal in the ischemic area based on six confocal images per brain. (d, e) Immunofluorescent staining for PDGFR*α* (green) show that the cell body area of PDGFR*α*^+^ NG2-glia cells (e, red) are increased in the ischemic border of the ipsilateral (ipsi) hemisphere 3 hours post MCAO in vehicle controls (two headed arrows) compared to the contralateral hemisphere and imatinib-treated mice (arrows). (f) Quantification of PDGFR*α*^+^ cell body area in the ischemic border based on ten confocal images per brain. (g) Immunofluorescence analysis of microglia activation was performed by staining for CD11b (green) on brain sections from imatinib-treated mice and vehicle controls at different times after MCAO. Microglia (arrows); activated microglia/macrophages (two headed arrows). (h) Quantification of activated microglia/macrophages was determined in the whole ischemic area in four sections per animal. ND; not detected. (i) qPCR analysis of RNA isolated from vascular fragments harvested 3 hours, 24 hours and 7 days post MCAO. *Il1a, Tnfa* and *Ccl2* are shown. Symbols represent individual data points and bars the group mean ± S.E.M. Cell nuclei were visualized by DAPI and endothelial cells by CD31. Statistical significance *p < 0.05; **p < 0.01; *** p < 0.001; in i) # refer to statistics between imatinib and vehicle and * to statistics between sham and MCAO (c, f, Student’s unpaired t-test; h, i, one-way ANOVA with Fisher’s LSD test). Representative maximum intensity projections of 11 µm (d); 8 µm (e); 12 µm (g) z-stacks are shown. Scale bars, 1000 µm (a); 50 µm (b); 25 µm (d); 10 µm (e, g).

### Imatinib reduces PDGFR*α* fibrotic scar formation through inhibition of myofibroblast transdifferentiation without affecting the formation of the astrocyte- or NG2-glia scar 7 days after MCAO

Reactive gliosis is an immediate and critical response to CNS injury, that is believed to orchestrate the subsequent formation of the glial scar. The glia scar is in turn believed to contain the damage and thereby improve outcome after injury, although, sustained gliosis has been reported to be deleterious to functional recovery^19^. Hence, the role of the scar is highly debated. To investigate whether the early effect of imatinib on reactive gliosis affects glial scar formation in the subacute/chronic tissue remodeling phase 7 days post MCAO, we performed immunofluorescent co-stainings of PDGFR*α* with markers of the glial scar; GFAP for the astroglia scar (Fig. 4a) and NG2 for the NG2-glia scar (Fig. 4b). These stainings revealed that imatinib treatment did not markedly affect the GFAP^+^ or the NG2^+^ scar, whereas it significantly reduced formation of a PDGFR*α*^+^ scar 7 days post MCAO compared to vehicle controls (Fig. 4c). We noticed that the PDGFR*α*^+^ scar was localized on the ischemic core side of the GFAP^+^ scar and that there was very little, if any, co-localization of GFAP and PDGFR*α* expression in the scar (Fig. 4a, higher magnification shown in Extended data Fig. 3). The PDGFR*α*^+^ scar appeared more disorganized in vehicle controls than in imatinib-treated animals and was embedded within the NG2^+^ scar (Fig. 4b). However, contrary to the normal expression pattern of PDGFR*α* in the brain (see Extended data Fig. 1), the majority of the parenchymal PDGFR*α* expression within the scar was detected in NG2^-^ cells (Fig. 4d). This ectopic expression of PDGFR*α* was found to co-localize with *de novo* parenchymal expression of ASMA (Fig. 5a – d), a commonly used marker for myofibroblast transdifferentiation/expansion following wound repair in peripheral tissue^38^. It should be noted that in the healthy murine brain ASMA expression is restricted to vSMC and no parenchymal expression of ASMA is observed (Extended data Fig. 4a). Analysis of the ASMA stainings revealed that MCAO induced pronounced myofibroblast transdifferentiation/ expansion in the PDGFR*α*^+^ portion of the scar of vehicle controls, which was much reduced by imatinib treatment (Fig. 5a, b). It appeared as if the double positive ASMA^+^ PDGFR*α*^+^ cells were leaving the vessel wall (Extended data Fig. 4b, c), suggesting the myofibroblasts might originate from perivascular cells, although this will require in depth lineage tracing analysis. Since myofibroblasts in peripheral organs are known to secrete extracellular matrix (ECM) molecules that further promotes disease pathogenesis^38^, we next performed stainings for the ECM glycoprotein fibronectin 7 days post MCAO (Fig. 5e,f).

**Figure 4.**
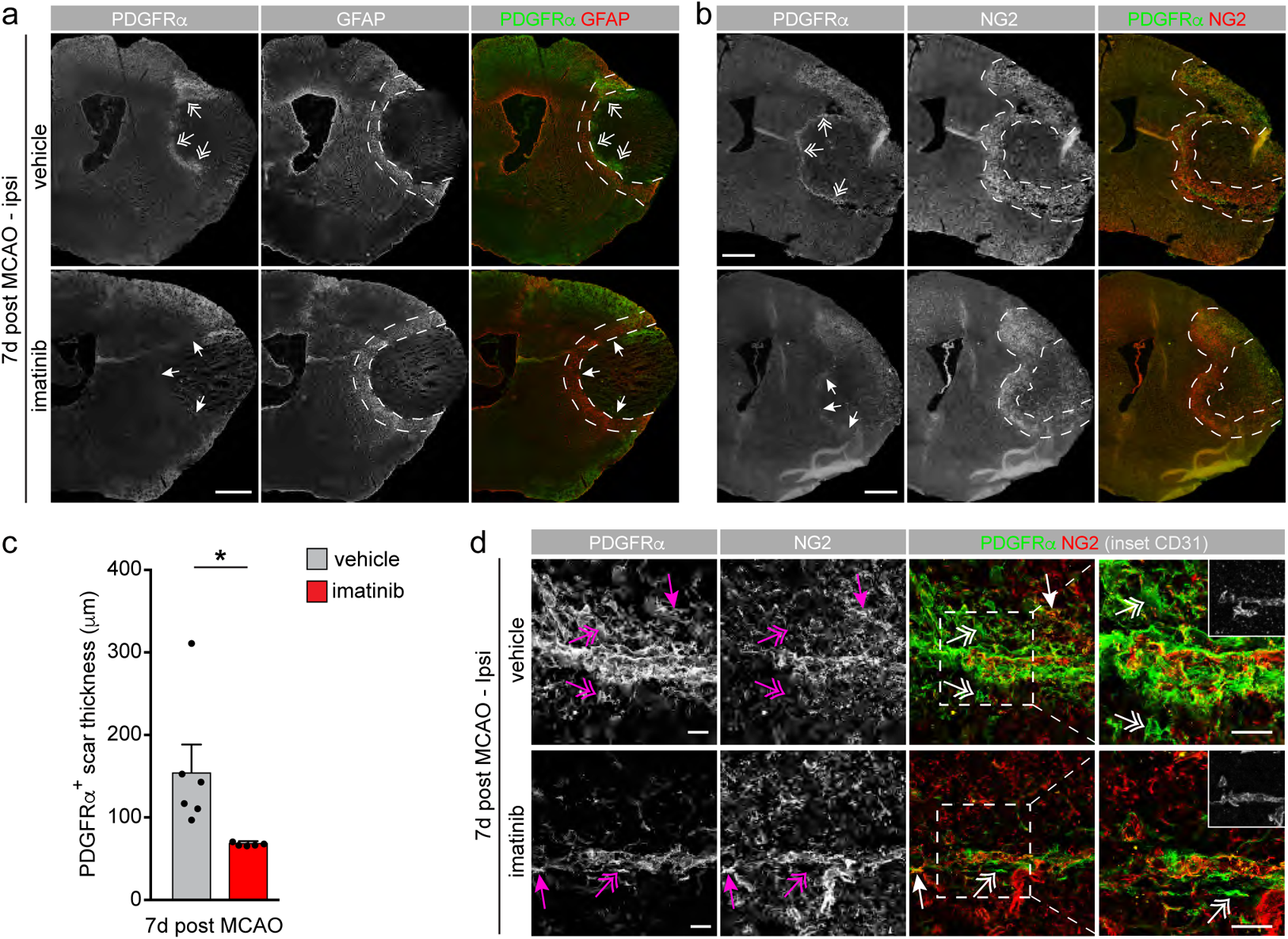
Imatinib controls the size of the PDGFR*α*^+^ scar 7 days after MCAO. Immunofluorescent co-stainings of PDGFR*α* with GFAP or NG2. (a – b) Stitched tiles of epifluorescent images (taken with a 10x objective). (c) Imatinib treatment reduced the formation of the PDGFR*α*^+^ scar 7 days post MCAO (arrows) compared to vehicle controls (two headed arrows), without affecting the thickness of the GFAP^+^ (a) or NG2^+^ (b) scar (demarcated with dashed lines). Quantification of PDGFR*α*^+^ scar thickness was determined by 20 individual measures along the scar. (d) Confocal images of the staining in (b) showing that MCAO induced ectopic expression of PDGFR*α* in NG2^-^ non-perivascular cells (two headed arrows) as opposed to normal expression in NG2^+^ glia cells (arrows). Symbols represent individual data points and bars the group mean ± S.E.M. Statistical significance *p < 0.05 relative to control (Student’s unpaired t-test). Representative maximum intensity projections of 11 µm z-stacks are shown in (d). Scale bars, 1000 µm (a, b); 25 µm (d).

**Figure 5.**
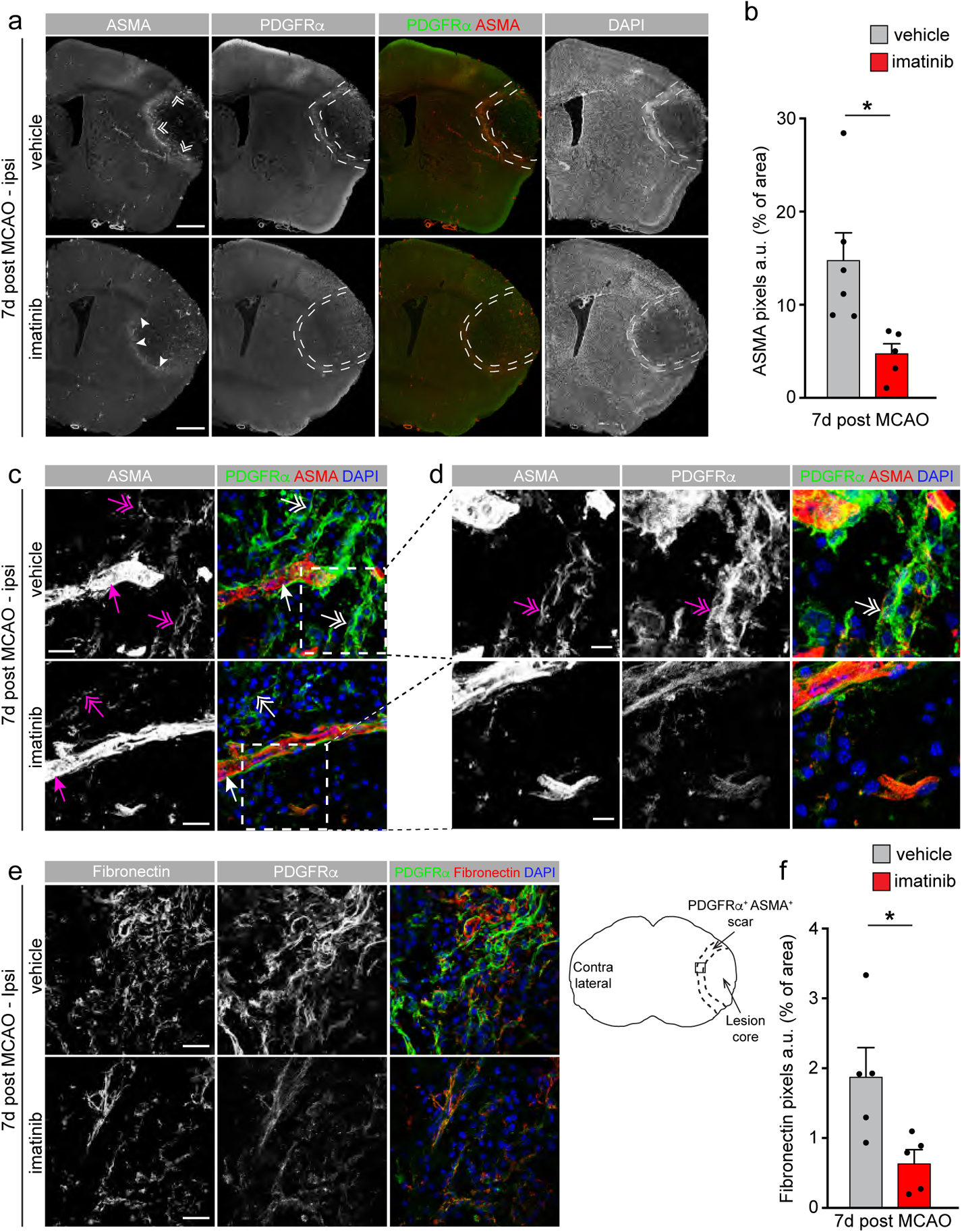
Imatinib reduces MCAO-induced myofibroblast expansion. Immunofluorescent co-stainings of PDGFR*α* with ASMA or fibronectin on brain sections from imatinib-treated and vehicle control mice 7 days post MCAO. (a) Stitched tiles of epifluorescent images (taken with a 10x objective). Imatinib treatment reduced ASMA^+^ signal in the scar (two headed arrowheads) compared to vehicle controls (arrowheads). The ectopic ASMA expression was localized to the highly nucleated PDGFR*α* scar (demarcated). (b) Quantification of ASMA signal in the entire PDGFR*α*^+^ highly nucleated scar demarcated in (a). (c, d) Confocal images show co-expression of ASMA with PDGFR*α* in parenchymal cells (two headed arrows) but not in vSMC (arrows). (d) Higher magnification of boxed area in c. (e) Confocal images of fibronectin and PDGFR*α* co-stainings taken in the highly nucleated PDGFR*α*^+^ scar region. (f) Quantification of fibronectin signal based on five confocal images per brain. Symbols represent individual data points and bars the group mean ± S.E.M. Statistical significance *p < 0.05 relative to control (Student’s unpaired t-test). Representative maximum intensity projections of 10 – 11 µm confocal z-stacks are shown. Scale bars, 1000 µm (a); 25 µm (c, e) and 10 µm (d).

These analyses reveled that the PDGFR*α*^+^ cells were surrounded by fibronectin-positive ECM and that less fibronectin-positive ECM was detected in the ASMA^+^ PDGFR*α*^+^ scar of imatinib-treated mice compared to controls, thus further supporting a role of imatinib in inhibiting myofibroblast transdifferentiation/expansion after MCAO. Interestingly it appeared as if the fibrotic scar within the lesion core was unaltered by imatinib treatment and that imatinib thus specifically targeted the expansion of the ASMA^+^ PDGFR*α*^+^ scar (Extended data Fig. 5). Because detrimental scarring is attributed to sustained myofibroblast expansion in pathologic wound healing in peripheral tissue, it is therefore possible that inhibition of this expansion is central to the beneficial effect seen when using imatinib in ischemic stroke.

### Imatinib progressively improves sensory-motor integration after MCAO

To test functional outcome following imatinib treatment we assessed lateralized sensory-motor integration 3 and 7 days after MCAO (Fig. 6a, study design) using the corridor task modified for mice^39, 40^. This test is based on the fact that mice with unilateral brain lesions display contralateral neglect and thus, preferentially explore/retrieve objects/food placed on the side ipsilateral to the lesion. Functional benefit of a treatment will thus result in reduced ipsilateral bias. We found that 3 days after MCAO all vehicle-treated mice preferentially explored sugar pellets from the side ipsilateral to the lesion (Fig. 6b). This ipsilateral exploration bias in vehicle controls largely remained 7 days after MCAO. However, imatinib treatment significantly reduced ipsilateral exploration bias compared to vehicle controls, both at 3 and 7 days post MCAO, although the effect was more pronounced at 7 days (Fig. 6b). In fact, at 7 days there was no significant difference between imatinib-treated mice and sham operated controls. As expected, imatinib significantly reduced lesion volume compared to vehicle controls 7 days post MCAO (Fig. 6c). Our analysis indicated a 37% reduction in lesion volume, which is in line with previously published data where therapeutic imatinib reduced the lesion by 34% 3 days post MCAO^7^. We found that lesion volume significantly correlated with exploration bias 7 days post MCAO, with all imatinib-treated animal clustering in the lower, left quartile whereas vehicle controls mainly clustered in the upper, right (Fig. 6d). This could suggest that functional outcome might merely reflect lesion size.

**Figure 6.**
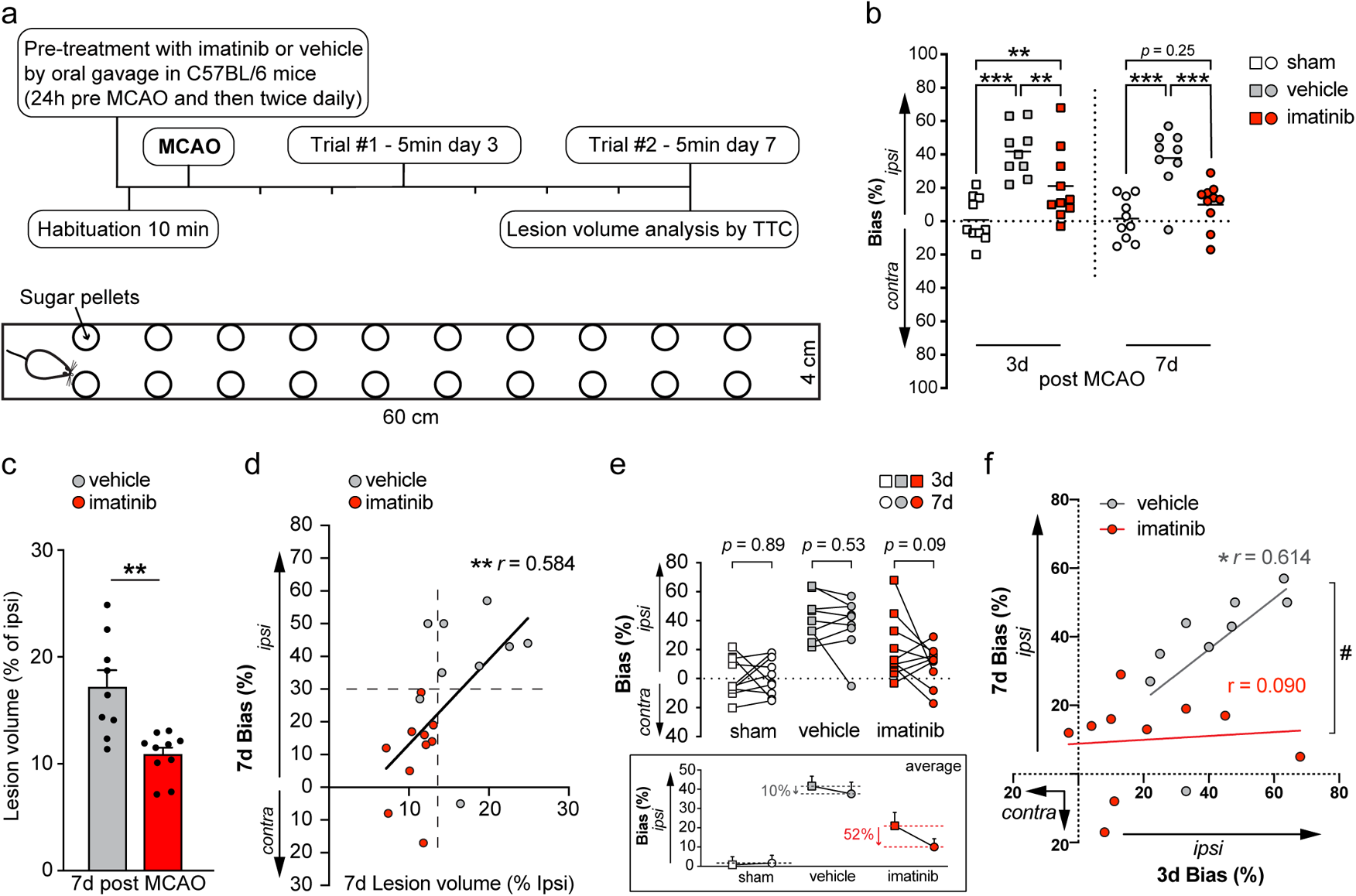
Imatinib progressively improves functional recovery after MCAO. (a) Schematic illustration of study design and the corridor task. After the mouse was placed at one end of the corridor, it was free to explore and eat sugar pellets at will. The number of pellet explorations made from either the left (ipsilateral to the lesion) or right (contralateral) side were counted. The trial was terminated when the mouse made 40 explorations or when the maximum trial time of 5 minutes was reached. Schematic corridor illustration is not to scale. (b) Exploration bias 3 and 7 days after MCAO in vehicle and imatinib-treated mice. MCAO led to a pronounced ipsilateral exploration bias in vehicle-treated mice at both time points, which was reduced by imatinib treatment. Sham operated mice were used as controls. (c) Quantification of lesion volume 7 days after MCAO in mice treated with either imatinib or vehicle. (d) Lesion volume significantly correlated with exploration bias at 7 days post MCAO, with all imatinib-treated mice clustering in the lower left quartile. (e) Change in exploration bias between day 3 and 7 post MCAO for each individual mouse. Inset shows group means ± S.E.M. and average reduction in ipsilateral bias. (f) Correlation of exploration bias 3 days post MCAO with the 7 day values in vehicle and imatinib-treated mice, respectively. Significant differences were detected between the lines #. Symbols represent individual data points and bars the group mean ± S.E.M. Statistical significance *p < 0.05; **p < 0.01; *** p < 0.001 (b, one-way ANOVA with Fisher’s LSD test; c, Student’s unpaired t-test; d, f, correlation and simple linear regression analysis; e, mixed-effects analysis with Fisher’s LSD test).

However, imatinib-treated mice displayed a greater reduction in ipsilateral exploration bias between day 3 and day 7 than vehicle controls (52% vs. 10% decrease, respectively) (individual values and group means shown in Fig. 6e), despite similarities in lesion volume reduction after imatinib treatment at the two different time points. Further, correlation of the 3-day bias data with the 7-day data showed a clear separation of the lines and significantly different correlations in vehicle vs. imatinib-treated mice (Fig. 6f). These data are consistent with the hypothesis that the consolidation of the fibrotic scar in imatinib-treated mice contributes to the improvement in functional outcome compared to the vehicle controls where the fibrotic scar is expanding. This suggests that the inhibition of myofibroblast expansion in the subacute/chronic phase of stroke recovery may enhance regenerative capacity and thereby contribute to improved functional recovery.

In conclusion, our findings show that imatinib dampens MCAO-induced cerebrovascular activation and the reactive gliosis response in the acute phase after ischemia. This is associated with reduced myofibroblast transdifferentiation and expansion in the subacute/chronic phase, which potentially improves regenerative capacity and thereby contributes to functional recovery in stroke management (Schematic illustration Fig. 7).

**Figure 7.**
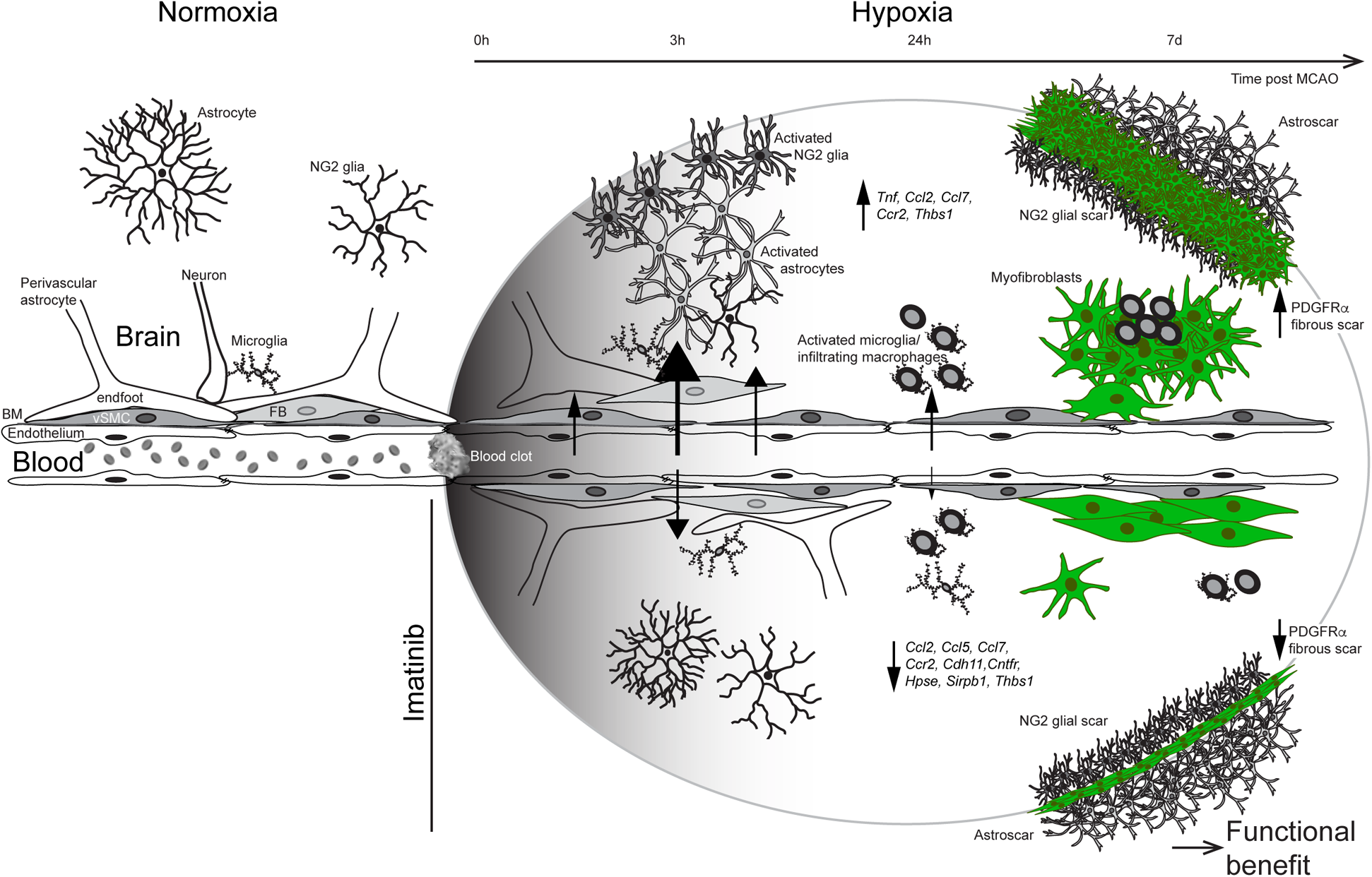
Schematic summary of imatinib-mediated effects post MCAO. To the left, the NVU during normoxia. The endothelium maintains a tight barrier with the support of the basement membrane (BM), vascular mural cells (including vascular smooth muscle cells, vSMC) and perivascular astrocytes and fibroblast like cells (FB). During normoxia non-vascular astrocytes, NG2 glia and microglia are ramified. To the right, the NVU and cellular responses during hypoxia after MCAO without (depicted above vessel) or with (depicted below vessel) imatinib treatment. In the acute phase (hours) after onset of hypoxia, MCAO induces reorganization in the NVU, associated with increased vascular permeability, as well as an early reactive gliosis response, including activation of astrocytes, NG2 glia and microgliosis/macrophage recruitment. This is dampened by imatinib treatment and is supported by BBB transcriptome analyses showing that imatinib treatment downregulate expression of proinflammatory and profibrotic transcripts including *Ccl2, Ccl5, Cdh11, Cntfr* and *Sirpb1*. In the tissue remodeling subacute/chronic phase after MCAO (days), imatinib reduces transdifferentiation/expansion of myofibroblasts and PDGFR*α* fibrotic scar formation. Taken together this results in a progressive improvement in functional outcome.

## Discussion

The molecular and cellular mechanisms of scar formation in the CNS are still poorly understood. Here, we found that imatinib dampened the early reactive gliosis response in the acute phase of ischemia and reduced myofibroblast transdifferentiation/expansion and fibrotic scarring in the subacute/chronic phase. We have previously postulated that the mechanism for the effect of imatinib in stroke, downstream of BBB preservation, potentially involves inhibition of immune cell infiltration^5, 31^. However, our analyses here point at a very limited effect of imatinib on peripheral leukocyte recruitment post MCAO, suggesting that it is unlikely that the attenuation of ischemia-induced systemic inflammatory processes account for a significant part of the beneficial outcome associated with imatinib treatment^5, 7^. This is in line with studies failing to show beneficial effects with immunomodulatory treatments in clinical settings of ischemic stroke^41^. Instead our data reveal that in addition to protecting the BBB, imatinib also modulates formation of the PDGFR*α*^+^ myofibroblast portion of the fibrotic scar following MCAO. We hypothesize this might be central to the beneficial effect seen on neurological and functional outcome, especially given that sustained myofibroblast expansion is well known to be detrimental in healing processes in many organs, although less is known about this process in the CNS^38^. This hypothesis is further supported by our findings that consolidation of the PDGFR*α*^+^ fibrotic scar in imatinib-treated mice seemingly works to discriminate behavioral outcome.

CNS injury repair has long been viewed as being markedly different from repair processes in other organs, mainly because of the poor regenerative capacity and immune-privileged nature of the CNS^17^. The insufficient regenerative capacity has largely been ascribed to the formation of a glial scar, in particular the astrocytic scar has been regarded as a barrier to CNS axon regrowth^20^. Recent studies, however, have shown that the presence of an astrocytic scar is not a principal cause for the failure of CNS axons to regrow. Instead, scar-forming astrocytes have been found to permit and support robust amounts of CNS axon regeneration in an experimental murine model of spinal cord injury^21^. These controversies have raised the question whether there might be different subtypes of reactive astrocytes eliciting different protective/harmful functions following injury and whether these potentially different astrocyte pools are activated in a time- and disease-specific manner. Toward this end, recent work has demonstrated that there are at least two different types of reactive astrocytes, neurotoxic A1 astrocytes and neuroprotective A2 astrocytes, which are induced in disease-specific manners^42, 43^. Our data support the existence of different subtypes of disease activated astrocytes. For example, we report a strong effect of imatinib on preserving perivascular astrocyte coverage in the NVU in the acute phase after MCAO, likely via direct inhibition of MCAO-induced PDGFR*α* signaling in these cells. Thus, supporting a protective role of these perivascular cells in maintaining vessel health and integrity. However, imatinib treatment also inhibited MCAO-induced activation of, what appears to be harmful, non-perivascular astrocytes in the ischemic core in the acute phase after MCAO induction. As these parenchymal astrocytes do not express PDGFR*α*, nor any of the other receptor tyrosine kinases known to be targeted by imatinib, including PDGFR*β*, Abl and c-Kit^14^, it seems likely this is mediated through an indirect mechanism. We speculate these parenchymal astrocytes might be activated by cytokines, including IL-1α, TNF*α* and C1q, which we found to be strongly upregulated in the vascular fragments after ischemia and normalized by imatinib, since these factors are known to stimulate astrocyte activation^44–46^ and potently induce activation of neurotoxic A1 astrocytes^43^. Interestingly though, the astrogliosis dampening effect of imatinib within hours following MCAO did not translate into a differential thickness of the GFAP^+^ astrocyte scar a week after injury. This might suggest that the astroglia scar originates from a subset of reactive astrocytes, not targeted by imatinib.

The formation of a scar following CNS injury, despite the potential inhibitory effect on axon regeneration, is thought to be crucial in order to seal off the lesion site from unaffected brain regions, thereby allowing temporal and spatial control of tissue remodeling of the injured tissue^17^. However, equally important as the formation of the scar is the termination of the scarring process. In most tissues this final key event of the normal healing process includes the appearance of injury-induced myofibroblasts contributing to ‘scar contraction’ and resolution of the injury by generation of strong contractile forces^17, 22^, although very little data is available on this process in CNS healing. In addition to their contractile role, myofibroblasts also secrete ECM molecules that influence mechanical signaling and cell adhesion. Persistence and/or dysregulation of ‘scar contraction’ and excessive ECM deposition leads to distortion of the parenchymal architecture, promoting disease pathogenesis and in worst cases tissue failure, a process referred to as fibrosis. Here we report that genes important for the fibrotic response, including *Ccl2, Il1a, Tnfa, Ccl5, Ccl7, Fgr,* and *Itgax*, are upregulated in vascular fragments post MCAO and subsequently downregulated by imatinib, suggesting that imatinib might control MCAO-induced fibrosis. This is of particular interest since the fibrotic component of the CNS scar has been largely overlooked and has only recently emerged as a potential target in various CNS lesions (reviewed in^47^). Here we show that the photothrombotic model of MCAO utilized in this study induces transdifferentiation/ expansion of myofibroblasts, as illustrated by *de novo* expression of ASMA in activated parenchymal cells, at the rim of the astroglia scar 7 days post MCAO. The ASMA^+^ myofibroblasts were found to co-express PDGFR*α*. It is possible that these cells arise from perivascular cells as has been suggested in wound healing processes in peripheral organs^22^ and that they upregulate ASMA as they transdifferentiate into contractile myofibroblasts leaving the vessel wall, although this will require further analysis e.g. by genetic lineage tracing. Imatinib treatment reduced this myofibroblast response, as seen in the immunohistochemical analyses, but also indicated in the gene expression analyses where genes highly implicated in activation and proliferation of myofibroblasts^48^, including *Pdgfra, Il1a, Il1r2, Ccl2, Ccl5,* and *Hpse*, were downregulated. Similar to the heterogeneity with injury-induced astrocytes, it has been suggested that the ASMA^+^ myofibroblasts only accounts for a subset of injury-induced mesenchymal cells^22^. Different subsets of activated mesenchymal cells have been proposed to vary spatiotemporally after injury as well as to elicit different protective/harmful functions in the repair processes. Since imatinib treatment resulted in a myofibroblast scar that appeared to be more well-structured than in the untreated controls and inhibited, but did not completely block, its formation, this might indicate differential effects on distinct subsets of activated mesenchymal cells. Understanding the cellular origin and potential multilineage differentiation of the injury-induced mesenchymal cells is therefore a central issue for future research.

The potential progenitors for myofibroblasts are a matter of some debate and have been proposed to include epithelial-/endothelial-mesenchymal transition; circulating bone marrow-derived fibrocytes; tissue-resident fibroblasts or other mesenchymal cells related to blood vessels, such as pericytes, adventitial cells, and mesenchymal stem cells^22^. The currently prevailing dogma however seems to lean toward the latter, where specific subsets of tissue-resident mesenchymal cells, mainly localized in a perivascular position, serve as the major source for ECM-producing cells after injury. Some of these perivascular progenitors have in peripheral organs been shown to express PDGFRα. For example, in the liver it has been shown that the primary progenitors for myofibroblasts are perivascular hepatic stellate cells and specific deletion of PDGFRα from these cells *in vivo* resulted in reduced myofibroblast transdifferentiation and fibrosis in a model of hepatotoxic liver injury^28^. In the CNS, lineage tracing analyses suggest that perivascular cells in the NVU are important for fibrotic scar tissue formation^49, 50^ and that reduction of this scarring response promotes recovery after spinal cord injury in mice^51^. These perivascular cells were shown to express PDGFRα and the authors referred to them as type A pericytes^49^, although this has later been challenged by single-cell analyses suggesting the perivascular PDGFRα^+^ cells are not pericytes and should instead be referred to as fibroblast-like cells^33^. We, on the other hand, have been referring to the perivascular PDGFRα^+^ cells in the CNS as perivascular astrocytes based on immunofluorescent studies indicating co-expression of GFAP and AQP4 with PDGFRα along medium to large size vessels throughout the CNS (Extended data Fig. 1 and 7,^8, 52^). It should however be noted that single-cell RNA sequencing of the NVU in mice failed to detect GFAP and AQP4 expression in the PDGFRα^+^ perivascular population^33^. Then again, GFAP was not readily detected in this dataset^33^ and further, cell type specific ablation of PDGFRα expression from perivascular cells in the CNS have been reported utilizing *GFAP-Cre* mice^8^. Whether the proposed names of the PDGFRα^+^ perivascular cells are referring to the same pool of cells or to diverse subpopulations of cells remain to be determined. However, the latter seems likely and is supported e.g. by the fact that type A pericytes are distributed along the capillary bed as well as upstream the arterial/venous vasculature^49, 50^, whereas PDGFRα^+^ perivascular cells are exclusively found around medium/large size vessels and are never distributed along capillaries.

Scar formation is not only fundamental in the healing process after stroke but also in other neurological pathologies such as multiple sclerosis (MS), Alzheimer’s disease or traumatic injury to the brain or spinal cord. A recent study demonstrated that a fibrotic scar forms in the spinal cord following immune infiltration during experimental autoimmune encephalomyelitis (EAE), the animal model of MS, and plays a role in regulating disease severity^53^. Interestingly, we have recently published that imatinib or anti-PDGF-CC treatment ameliorates EAE severity by blocking the loss of BBB integrity^12, 13^. Data presented here point to an additional mechanism for imatinib / anti-PDGF-CC mediated amelioration of neuroinflammation, namely reduced fibrosis and scar contraction.

In conclusion, our data suggest that imatinib improves outcome of ischemic stroke by inhibiting disease-induced cerebrovascular changes and thereby expression of profibrotic genes from the activated BBB in the acute phase after ischemia onset. Consequently, this leads to reduced myofibroblast transdifferentiation/expansion in the subacute/chronic phase. This order of events is supported by our findings that imatinib appeared to be more beneficial if administered to patients within 5 hours from stroke onset, coinciding with the timing of the highest peak of cerebrovascular permeability, whereas the beneficial effect tailed off if imatinib treatment was initiated at a later time point^5^. We propose the effect is mediated through inhibition of ischemia-induced activation of PDGFRα signalling in perivascular cells in the NVU, which is in line with our previous findings showing that PDGFRα phosphorylation is induced in the NVU within hours following MCAO induction^31^. It should be noted though that imatinib is not exclusively inhibiting PDGFRα but also PDGFR*β*, Abl and c-kit, of which PDGFR*β* is of particular interest given that this receptor is also expressed in perivascular PDGFRα^+^ cells of the NVU^33, 49^ and has been implicated in myofibroblast activation in injury responses in peripheral organs^54^. Contrary to PDGFRα however, PDGFR*β* is also expressed in vascular mural cells, both in pericytes distributed along the capillaries and vSMC^33^. Also, during ischemic stroke, expression of PDGFR*β* is highly upregulated throughout the ischemic core region, and not selectively only at the rim of the astroglia scar^50, 55^. More importantly, reduced ischemia-induced expression of PDGFR*β* in the lesion core has been found to correlate with an enlargement of infarct volume after MCAO^56^ and stromal PDGFR*β*^+^ pericytes have been reported not to contribute to ECM production in the fibrotic lesion after stroke^57^. The latter is however in stark contrast to the lineage tracing studies of type A pericytes^50^, where type A pericytes were shown to give rise to nearly the entire fibrotic scar in the lesion core after cortical MCAO, a region of the fibrotic scar that appeared unaffected by imatinib treatment in our analyses. Nevertheless, this highlights a heterogeneity of the fibrotic scar, where it seems that reducing the PDGFRα^+^ myofibroblast portion is beneficial, whereas tampering with the fibrotic core portion of the scar might be detrimental, although this requires further investigation. In addition, it appears the different parts of the fibrotic scar stem from discrete progenitors, thus opening the possibility of differential targeting. Further studies, e.g. utilizing more specific ways to explicitly target the PDGFRα pathway genetically or pharmacologically, including the use of a monoclonal PDGF-C antibody^58, 59^, as well as longitudinal *in vivo* imaging approaches to lineage trace the response of these enigmatic perivascular PDGFRα^+^ cells to ischemia, are therefore warranted. Taken together, this offers new opportunities to study and understand the multidimensional roles and complex cellular interactions of fibrotic scar formation and function in ischemic stroke but potentially also in other neurological pathologies.

## Online materials and Methods

### Ethical statement

All experiments in this study were approved and performed in accordance with the guidelines from the Swedish National Board for Laboratory Animals and the European Community Council Directive (86/609/EEC) and were approved by the North Stockholm Animal Ethics Committee and the Institutional Animal Care and Use Committee of Unit for Laboratory Animal Medicine at the University of Michigan, respectively.

### Mouse model of ischemic stroke

Age- and gender matched C57BL/6 wild type mice were anesthetized with isoflurane and securely placed under a dissecting microscope. The left middle cerebral artery (MCA) was exposed and a laser Doppler flow probe (Type N (18 gauge), Transonic Systems) was placed on the surface of the cerebral cortex located 1.5 mm dorsal median from the bifurcation of the MCA as described before^7^. The probe was connected to a flowmeter (Transonic model BLF22) and relative tissue perfusion units (TPU) data was recorded with a continuous data acquisition program (Windaq, DATAQ Instruments). The photoactivatable dye Rose Bengal (Fisher Scientific) was diluted to 10 mg/ml in phosphate buffered saline (PBS) and then injected intravenously with the final dose of 40 mg/kg. A 3.5-mW green light laser (540 nm, Melles Griot) was directed at the MCA from a distance of 6 cm at the onset of the injection, and the TPU of the cerebral cortex was recorded. Total MCA occlusion (MCAO) was achieved when the TPU dropped to less than 30% of pre-occlusion levels and to achieve a stable clot the laser was left on for 20 minutes. As controls, C57BL/6 mice from the same cohort were shaved, skin was cut, the muscle retracted and either laser or Rose Bengal was used (sham operated mice). To alleviate postoperative pain, carprofen (Rimadyl Vet.®, Pfizer) was administered by subcutaneous injection at the start of the surgical procedure.

### Imatinib treatment

To block PDGFRα activation after stroke, mice were treated with the tyrosine kinase inhibitor imatinib (Novartis, Switzerland alternatively Mylan AB, Sweden) three times (morning-night-morning) before MCAO (daily dose 250 mg/kg by oral gavage). For harvesting at 24 hours post MCAO the mice were additionally gavaged once post MCAO and the mice sacrificed 7 days post MCAO were gavaged every 12 hours until the end of the experiment. As control, mice were orally gavaged the corresponding times with vehicle (H_2_O for the functional corridor tests, PBS for all other experiments). Imatinib tablets were crushed into a fine powder, solubilized in vehicle, vortexed and incubated at 37°C (water bath) for 5 minutes. Insoluble components were spun down in a table microcentrifuge (16,100 x *g*) for 10 minutes. The supernatant was used for oral gavage performed with a steel gavage needle for mice.

### Evans blue dye (EB) extravasation

For analysis of cerebrovascular permeability after MCAO, stroked mice were injected with 100µl of 4% Evans blue dye (Sigma-Aldrich) intravenously 1 hour prior to sacrifice. The animals were then transcardially perfused with PBS for 5 minutes under isoflurane anesthesia and the brains removed and photographed using a Canon PowerShot SX200IS camera.

Thereafter the brains were separated into hemispheres and each hemisphere was then homogenized in N,N-dimethylformamide (Sigma-Aldrich) in Precellys lysing tubes and centrifuged twice at 16,100 x *g* for 20 minutes. The supernatants were collected, EB extravasation determined by absorbance measurement and quantified separately in the ipsi- and contralateral hemispheres as described in^60^. Background EB level in the non-ischemic contralateral hemisphere was subtracted from the ischemic hemisphere. EB levels in each hemisphere were calculated using the following formula:

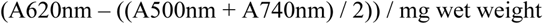

### Immunofluorescence and confocal microscopy

Tissue preparation for sectioning and immunostaining was conducted using standard protocols. Mice were anesthetized with isoflurane and transcardially perfused with PBS followed by fixation with 4% paraformaldehyde (PFA) in PBS. The brains were dissected and postfixed in 4% PFA 1 hour at room temperature (RT) and then kept in 30% sucrose (cryopreserved) 4°C overnight. The brains were thereafter either cut in 50 µm vibratome sections or cryopreserved in OCT and sectioned in a cryostat (12-14 μm coronal brain sections). The cryosections were permeabilized and then blocked with DAKO blocking solution, followed by incubation over night at 4°C with primary antibodies diluted in blocking solution. Vibratome sections were permeabilized and blocked with 1% bovine serum albumin in 0.5% TritonX-100/PBS and stained free floating in 24-well plates. The specific primary antibodies used were: goat anti-mouse PDGFR*α* (AF1062, R&D systems), rabbit anti-mouse GFAP (Z0334, DAKO), rat anti-GFAP (clone 2.2B10, 13-0300, Invitrogen), rabbit anti-rat NG2 (AB5320, Merck/Millipore), mouse anti-*α* Smooth Muscle Actin-Cy3 (ASMA, C6198, Sigma-Aldrich), goat anti-mouse CD31 (AF3628, R&D systems), rat anti-mouse CD31 (553370, BD Pharmingen), rat anti-mouse CD11b (550282, BD Pharmingen), rabbit anti-mouse CD3 (C7930, Sigma-Aldrich), rabbit anti-mouse MPO (ab208670, Abcam), rat anti-mouse B220 (CD45R, MAB1217, R&D systems), rabbit anti-fibronectin (F3648, Sigma-Aldrich) and rabbit anti-aquaporin 4 (AB2218, Merck/Millipore). Secondary antibodies (donkey anti-rat, donkey anti-goat and donkey anti-rabbit) conjugated with Alexa Fluor dyes were obtained from Invitrogen. DAPI (4′,6-Diamidino-2-Phenylindole, Dihydrochloride, 0.2 μg/ml) was included in the last PBS wash to visualize the nuclei. Following immunofluorescent staining the sections were mounted using ProLong Gold Antifade reagent (Molecular Probes). All images were acquired at RT with a Zeiss LSM700 confocal microscope or a Zeiss Axio Observer Z1 inverted microscope and the ZEN 2009 software (Carl Zeiss Microimaging GmbH). Stained brain sections from imatinib-treated mice or vehicle controls 3 hours, 24 hours and 7 days post MCAO were analyzed by two independent investigators blinded to the study group.

For all the quantifications of antibody immunoreactivity done in this study, images were acquired using the same settings (within the respective staining experiment) and taken in comparable anatomic positions for each animal. For quantification of GFAP^+^ and PDGFR*α*^+^ vessels in the ischemic area 3 hours post MCAO, eight confocal z-stacks (8 μm) were acquired per brain (n = 3 animals per treatment). The vessels were counted manually from the maximum intensity projection image and average number of positive vessels per mm^2^ was determined for each animal. For quantification of astrogliosis 3 hours post MCAO, six single-plane confocal images per brain were acquired in the ischemic area (n = 3 – 4 animals per treatment. Confirmed in a separate cohort and staining experiment using n = 3 animals per treatment). The area of antibody immunoreactivity above a set threshold was determined using ImageJ64. For quantification of PDGFR*α*^+^ cell body size in the ischemic border 3 hours post MCAO, ten confocal z-stacks (11 μm) were acquired per brain and average cell size determined from the maximum intensity projection image using the Volocity image software (n = 3 – 4 animals per treatment. Confirmed in a separate cohort and staining experiment using n = 3 animals per treatment). For comparison, images were also acquired and average cell size determined in the contralateral hemispheres. For quantification of immune cell infiltration at different times post MCAO, immune cells were counted manually and average number of positive cells per mm^2^ was determined in 4 different fields of view within the ischemic area per animal (n = 3 – 4 animals per treatment and time point). For quantification of PDGFR*α*^+^ scar thickness 7 days post MCAO, four confocal z-stacks (10 μm) were acquired covering the entire scar in each brain (n = 5 – 6 animals per treatment. Confirmed in a separate cohort and staining experiment using n = 4 animals per treatment). The average scar thickness was determined from 20 different positions along the scar (5 per image) for each animal. For quantification of ASMA expression 7 days post MCAO, immunoreactivity above a set threshold was determined in the PDGFR*α*^+^ scar region from the stitched tiled epifluorescent images using Image J.

All images shown are representative of the respective staining and were processed and analyzed using Volocity 3D image analysis software (PerkinElmer, Waltham, MA, USA), Adobe Photoshop CC (Adobe, San Jose, CA, USA) or Image J (National Institutes of Health, Bethesda, MD, USA). The result from all the fields of view in a given animal was averaged to obtain the value for that individual. Individual values and group mean ± S.E.M. are shown.

### Isolation of cerebrovascular fragments and generation of mRNA

Cerebrovascular fragments were isolated from mice 3 hours, 24 hours or 7 days post MCAO or from sham operated mice. Mice were anesthetized with isoflurane and thereafter perfused with HBSS. The brains were rapidly dissected, placed in ice-cold HBSS and mechanically and enzymatically dissociated. Biotin rat anti-mouse CD31 antibodies (BD Biosciences) coupled to magnetic Dynabeads (Dynabeads biotin binder, Invitrogen) were used to pull out vascular fragments as described previously^61^. Total RNA was extracted from both the wash and the eluate fractions using the RNeasy kit (Qiagen, Hilden, Germany) and the QIAcube (Qiagen) including on column DNA-digestion for fully automated sample preparation. RNA concentration and purity were determined through measurement of A260/A280 ratios with a NanoDrop ND-1000 Spectrophotometer (NanoDrop Technologies, Wilmington, DE, USA). Confirmation of RNA quality was assessed using the Agilent 2100 Bioanalyzer (Agilent Technologies, Santa Clara, CA, USA). mRNA was subsequently either used for expression array analysis or cDNA generation for qPCR analysis. cDNA was prepared using the iScript kit (Biorrad, Hercules, CA, USA). The purity of the cerebrovascular fragments and the fraction of various cell types in the preparations was analyzed by real time quantitative PCR (qPCR) using primers for endothelial cells (*Cldn5*, *Pecam1*), vascular mural cells (*Pdgfrb*), astrocyte endfeet (*Aqp4*), and neurons (*Dlg4*). The analysis revealed high enrichment of endothelial cells (187 ± 29 fold for *Cldn5* and 78 ± 16 fold for *Pecam1*), followed by vascular mural cells (25 ± 6 fold) and astrocyte endfeet (13 ± 2 fold). The cerebrovascular fragments were devoid of neurons (0.1 ± 0.05 fold). The values represent the enrichment factor (± S.E.M.) in the vascular fragment eluates compared to the wash fraction (rest of the brain tissue). These results indicate high purity of the cerebrovascular fragments.

### Real-time PCR analysis

Real-time quantitative PCR was performed using KAPA SYBR FAST qPCR Kit Master Mix (2x) Universal (KAPA Biosystems) in Rotor-Gene Q (Qiagen) Real-time PCR thermal cycler according to the manufacturer’s instructions. Expression levels were normalized to the expression of *Rpl19*.

The following primers were used:

5’-GGTGACCTGGATGAGAAGGA, 5’-TTCAAGCTTGTGGATGTGCTC. (*Rpl19*); 5’-CTGTAGCCCACGTCGTAGC, 5’-ACAAGGTACAACCCATCGGC (*Tnfa*); 5’-ATCAGCAACGTCAAGCAACG, 5’-AAGGTGCTGATCTGGGTTGG (*Il1a*); 5’-TTAAAAACCTGGATCGGAACCAA, 5’-GCATTAGCTTCAGATTTACGGGT (*Ccl2*).

The following primers were used for confirmation of purity of the vascular fragments: 5’-TACTGGGCTTCGAGAGCATT, 5’-AGAGACGGTCTTGTCGCAGT (*Pecam1*); 5’-CACCTTCTCCAGTGTGCTGA, 5’-GGAGTCCATAGGGAGGAAGC (*Pdgfrb*); 5’-CGCCCCCTCTGGAACACAGC, 5’-TGCTGGAGGGCGAAGAAAACCG (*Dlg4*); 5’-ATGGTGGATCCCACACCGAG, 5’-AGGCGGTGGGGTAAGTGTG (*Aqp4*); 5’-GTGGAACGCTCAGATTTCAT, 5’-TGGACATTAAGGCAGCATCT (*Cldn5*).

Measurements are depicted as mean ± S.E.M. from the number of samples stated in the figure legends.

### Microarray and data analysis

10 nanograms of total RNA from each sample were used to generate amplified and biotinylated sense transcript cDNA from the entire expressed genome according to the Nugen Technologies, Inc. protocols Ovation® Pico WTA System V2 (M01224v2) and Encore Biotine Module (M01111v5). GeneChip® ST Arrays (GeneChip® Mouse Gene 2.0 ST Array) were hybridized for 16 hours in a 45°C incubator, rotated at 60 rpm. According to the GeneChip® Expression Wash, Stain and Scan Manual (PN 702731 Rev 3, Affymetrix Inc., Santa Clara, CA) the arrays were then washed and stained using the Fluidics Station 450 and finally scanned using the GeneChip® Scanner 3000 7G. The raw data were normalized in the free software Expression Console provided by Affymetrix (http://www.affymetrix.com) using the robust multi-array average (RMA) method first suggested by Li and Wong.

Subsequent analysis of the gene expression data was carried out in the freely available statistical computing language R (http://www.r-project.org) using packages available from the Bioconductor project (www.bioconductor.org). In order to search for the differentially expressed genes between the different groups an empirical Bayes moderated t-test was then applied, using the ‘limma’ package. To address the problem with multiple testing, the *P-*values were adjusted using the method of Benjamini and Hochberg.

To compare gene expression in vascular fragments from imatinib-treated mice versus vehicle controls 3 hours and 24 hours post MCAO, molecules from the dataset that met the < or > log2 (0.5) fold change and *P*-value < 0.05 cutoff were uploaded to the ingenuity pathways analysis platform (Ingenuity Systems, CA, USA, www.ingenuity.com). The molecules in this dataset were grouped in biological functions and/or diseases or were associated with a canonical pathway in Ingenuity’s knowledge base. Right-tailed Fisher’s exact test was used to calculate a *P*-value determining the probability that each biological function and/or disease assigned to that data set is due to chance alone. The significance of the association between the data set and the canonical pathway was measured in 2 ways: 1) A ratio of the number of molecules from the data set that map to the pathway divided by the total number of molecules that map to the canonical pathway is displayed. 2) Fisher’s exact test was used to calculate a *P*-value determining the probability that the association between the genes in the dataset and canonical pathway is by chance alone. Raw data are deposited on the NCBI Gene Expression Omnibus database (accession no. GSE137534). To compare our dataset to the harmonizome database^32^ we used the harmonizome dataset named “fibrosis, CTD Gene-Disease Associations”. In the harmonizome database every gene included in a certain functional annotation has a standardized value indicating the relative strength of the association.

Standardized values range from 1 to 2,88. We compared our dataset with all genes from the fibrosis dataset showing a higher standardized value than 1,5. The association with inflammation and metabolism was done manually and according to the current scientific knowledge.

### Corridor task and stroke volume

To assess functional recovery we measured lateralized sensory-motor integration using a corridor task adapted from ^39, 40^. Briefly, as the testing corridor a plexiglass box (L=60 cm x W=4 cm x H=15 cm) was used, where ten pairs of adjacent Eppendorf caps, each containing 4-5 sugar pellets (20 mg per pellet; TestDiet), had been placed at 5-cm intervals. A corridor with the same dimensions but without adjacent Eppendorf caps was used as the habituation corridor. 24 hours before MCAO, imatinib or vehicle-treated mice were habituated to the corridor by scattering sugar pellets along the corridor floor and allowing them to freely explore for 10 minutes. Sham operated mice were used as controls. Lateralized sensory-motor integration was tested 3 and 7 days after MCAO. On the testing day, mice were placed in the habituation corridor for 5 minutes in the absence of sugar pellets, then mice were transferred to one end of the testing corridor containing sugar pellets and video recorded for 5 minutes. All video recordings were analyzed by an investigator blinded to the experiment. A second investigator analyzed randomly selected videos independently to confirm scoring by the first investigator (approx. 16% of all videos were analyzed by two investigators). The number of ipsilateral and contralateral explorations relative to the stroked hemisphere were counted until the mouse made a total of 40 explorations or the video ended. An exploration was defined as a nose-poke into an Eppendorf cap, whether the sugar pellet was poked or eaten, and a new exploration was only counted by exploring a new cap. Data is expressed as ipsilateral bias, calculated as:

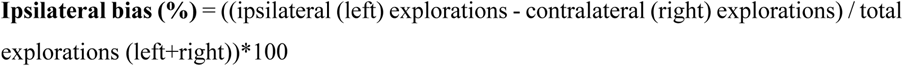

Following the last day of functional testing stroke volume analysis was performed as described previously^7, 31^. Briefly, brains were removed 7 days after MCAO and 2 mm thick coronal sections of the whole brain were stained with 4% 2,3,5-triphenyltetrazolium chloride (TTC) in PBS for 20 minutes at 37°C and then fixed in 4% paraformaldehyde solution for 10 minutes. TTC stained sections were captured with an Olympus digital C-3030 color camera attached to an SZ-60 Olympus microscope. The sections were analyzed with NIH Image J software using the following formula:

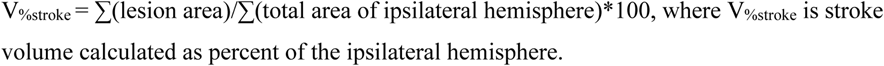

### Statistics

Data analysis was performed using GraphPad Prism 9 statistical software (GraphPad Software, La Jolla, CA, USA). For statistical analysis in any experiment with only two groups a two-tailed t-test was used. For experiments with more than two groups, statistical evaluation was performed using one-way ANOVA with Fisheŕs LSD with statistical significance defined as \**P* ≤ 0.05, \*\**P* ≤ 0.01 and \*\*\**P* ≤ 0.001. Data is represented as mean values ± S.E.M.

## Acknowledgements

We would like to thank the Array and Analysis Facility, Science for Life Laboratory at Uppsala Biomedical Center (BMC), Husargatan 3, 751 23 Uppsala; Yihang Li and Nadine Winkler for help with qPCR and stainings. This work was supported by the Karolinska Institutet (M.Z., M.Z.A., U.E., I.N., L.F.), the Swedish Research Council (Vetenskapsrådet) (L.F., 524-2010-7045; 521-2012-1853)(U.E., 2016-02593; 2017-01794) (I.N., FS-2008-90), the Swedish Governmental Agency for Innovation Systems (VINNOVA) (L.F., 2011-03503), Hållsten foundation (U.E., I.N., L.F.), Swedish Brain Foundation (U.E., I.N., L.F. FO2020-0037; FO2021-0039), Torsten Söderberg Foundation (U.E., M137/16), Neurofonden (M.Z., M.Z.A., S.A.L., I.N.), Olle Engkvist Byggmästare Foundation (S.A.L., SLS-499431), Ulla-Carin Lindquists foundation for ALS research (S.A.L.), Åhlen foundation (S.A.L.), the Royal Swedish Academy of Sciences (I.N.), Magnus Bergvalĺs foundation (I.N.), the Swedish Stroke Association/STROKE Riksförbundet (I.N.), and the Swedish Heart and Lung foundation (Hjärt-lungfonden)(U.E., I.N., 20150547).

## Compliance with ethical standards

Yes. All applicable international, national, and/or institutional guidelines for the care and use of animals were followed. All procedures performed in studies involving animals were in accordance with the ethical standards of the institution at which the studies were conducted.

## Ethical approval

All animal experiments were reviewed and approved by the local animal ethics committees.

## Data availability

Raw data are deposited on the NCBI Gene Expression Omnibus database (accession no. GSE137534).

