## Supplemental information for "Reduced myofibroblast transdifferentiation and fibrotic scarring in ischemic stroke after imatinib treatment"

Extended data Fig. 1. Zeitelhofer M et al.

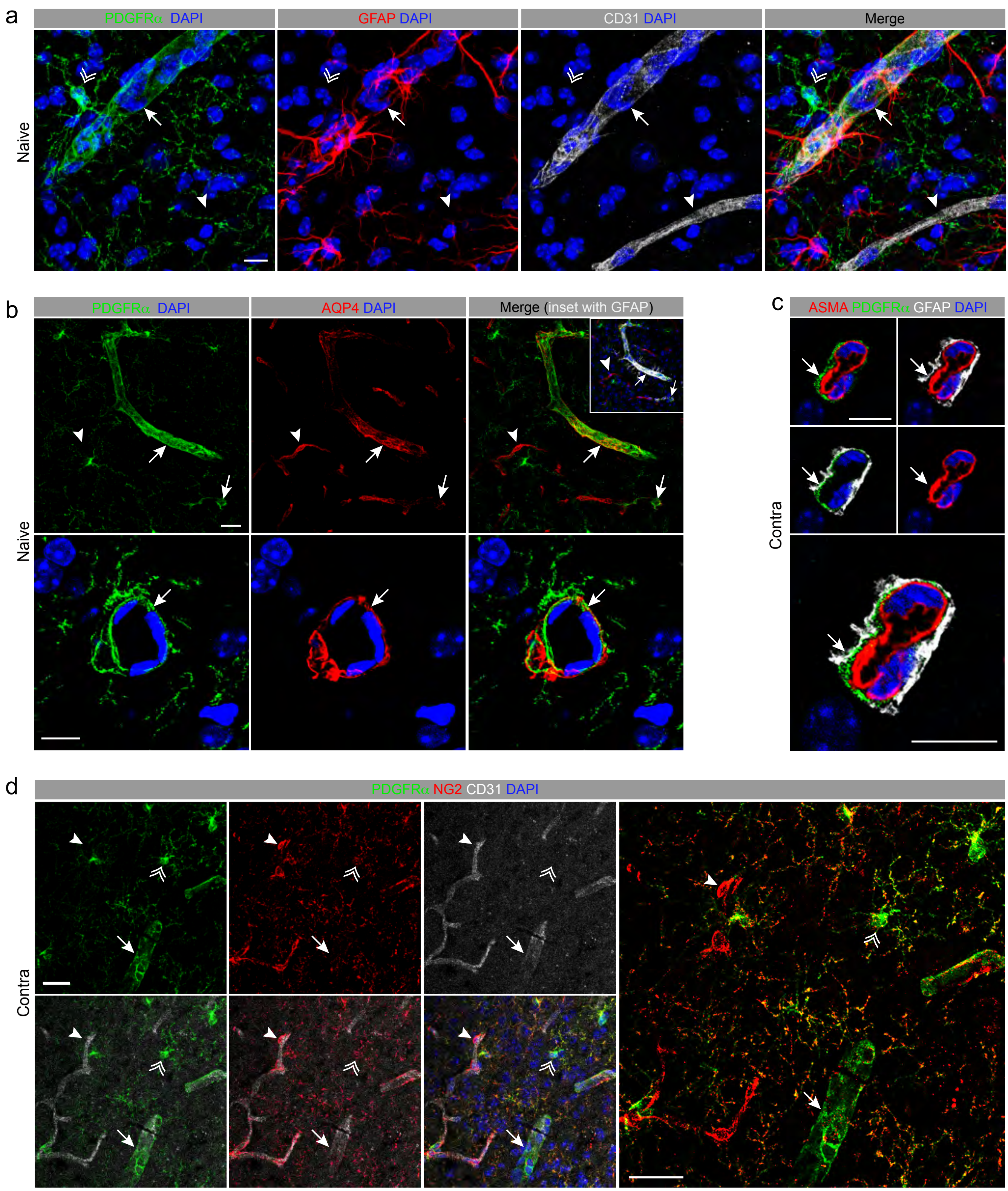

Extended data Fig. 2. Zeitelhofer M et al.

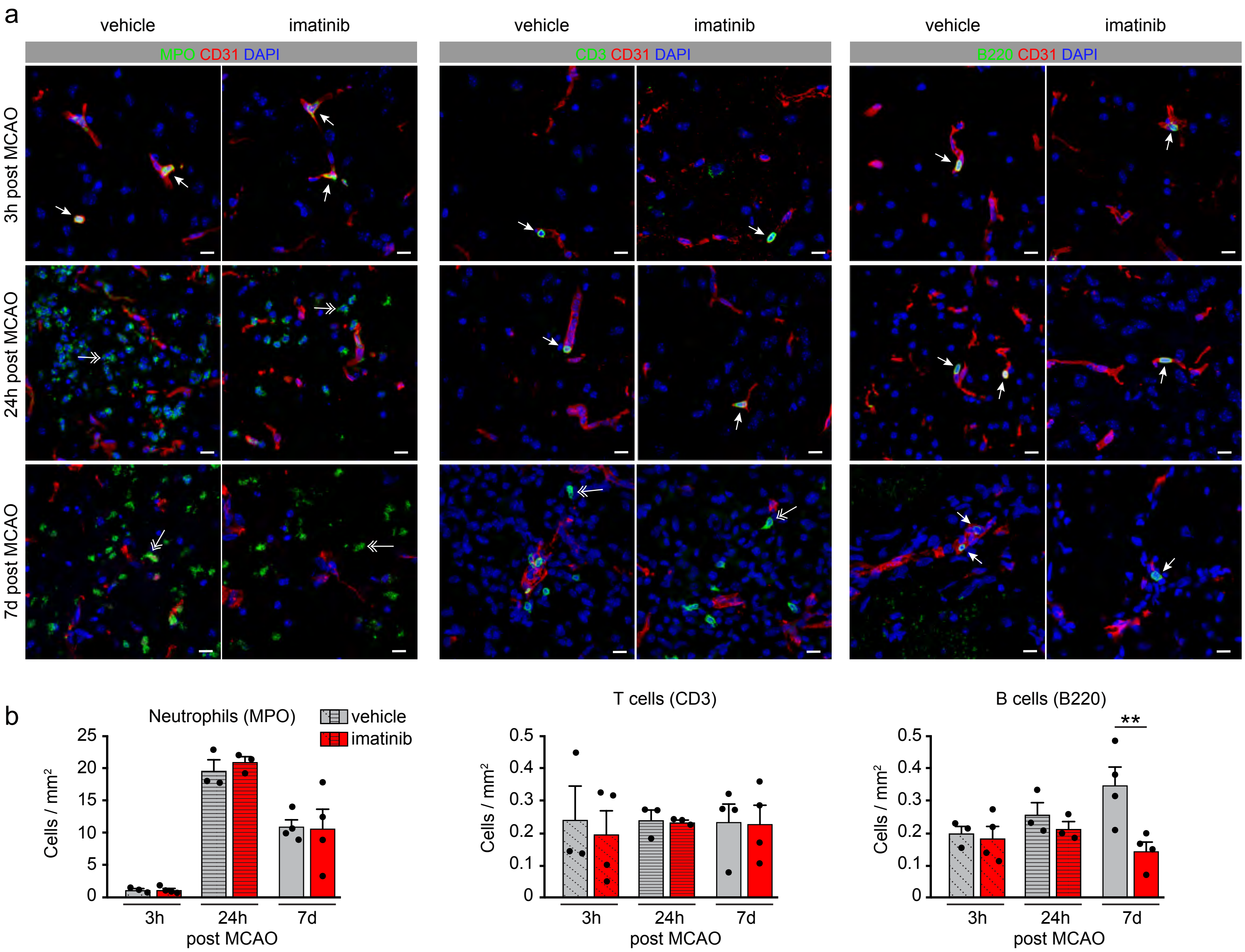

Extended data Fig. 3. Zeitelhofer M et al.

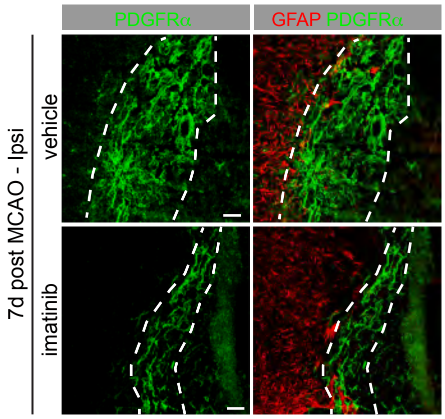

Extended data Fig. 4. Zeitelhofer M et al.

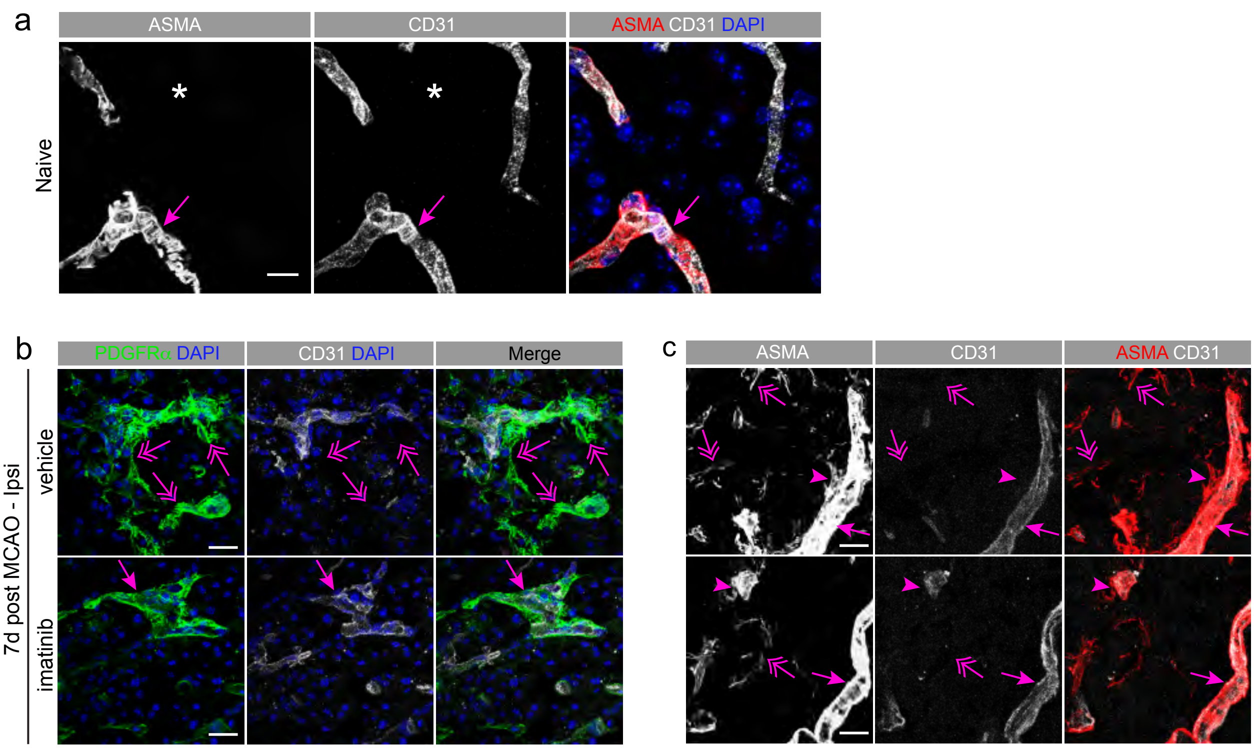

Extended data Fig. 5. Zeitelhofer M et al.

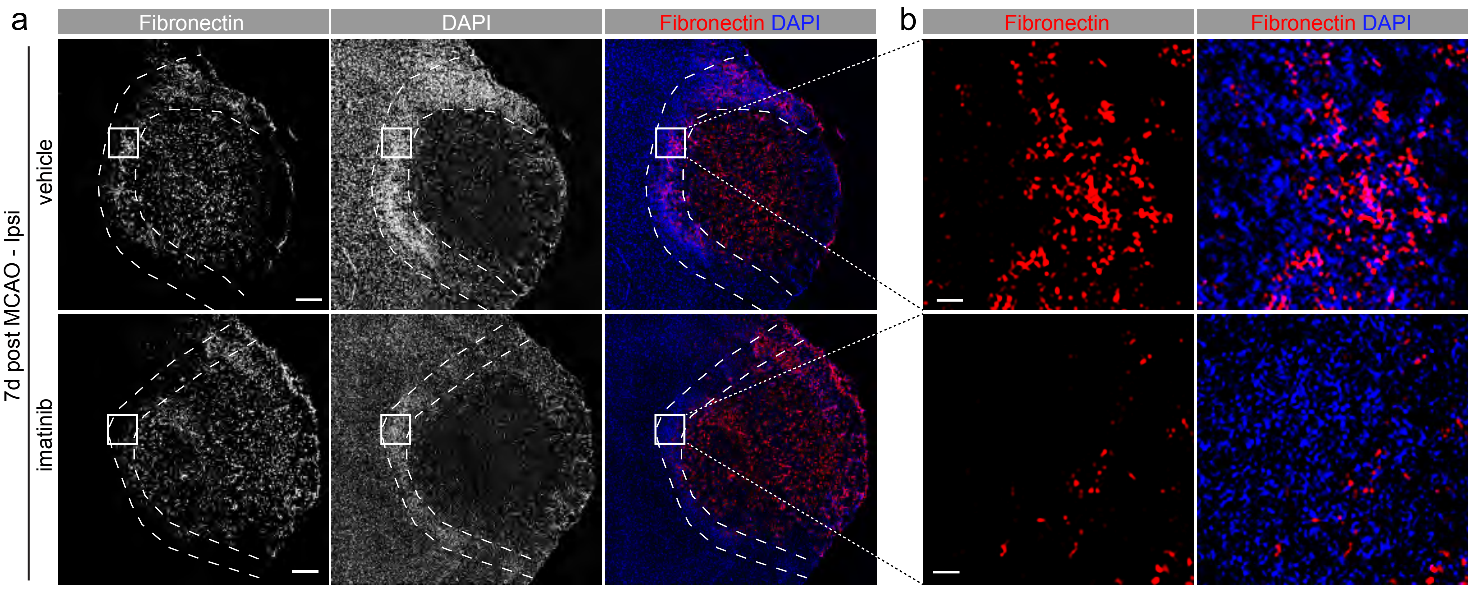

**Extended data Figure 1. Cellular organization in the unchallenged NVU.**

(a) Immunofluorescent co-staining for PDGFRα (green), GFAP (red), and the endothelial cell marker CD31 (white) in naïve wild type brain sections (vibratome free float stain). Perivascular PDGFRα is expresses around $GFAP^{+}$ vessels (arrows) but not around $GFAP^{-}$ capillaries (arrowheads). PDGFRα is also expressed in non-vascular parenchymal cells (two headed arrowhead). (b) Immunofluorescent co-staining for PDGFRα (green) and AQP4 (red) in naïve wild type brain sections (vibratome free float stain). Perivascular PDGFRα co-localize with AQP4 in $GFAP^{+}$ vessels (arrows, inset GFAP in white) but not in $AQP4^{+}$ $GFAP^{-}$ capillaries (arrowheads). (c) Immunofluorescent co-staining for PDGFRα (green) and GFAP (white) with the vascular smooth muscle cell (vSMC) marker ASMA (red) shows that double positive $PDGFR\alpha^{+}$;$GFAP^{+}$ cells are localized on the parenchymal side of the vessel wall, outside the vSMC lining in the contralateral hemisphere (arrows). (d) Immunofluorescent co-staining of PDGFRα (green), the pericyte/NG2-glia marker neuron glia antigen-2/CSPG4 (red), and the endothelial cell marker CD31 (white) in the contralateral hemisphere. PDGFRα in the CNS display high expression in NG2-expressing non-vascular parenchymal glia cells (two headed arrowhead), in addition to its expression in perivascular cells (arrow). NG2 expression is also detected in vascular mural cells (pericytes and vascular smooth muscle cells) (arrowhead). Cell nuclei were visualized by DAPI (blue). Representative maximum intensity projections of 18 μm (a, upper panel b), 3 μm (lower panel b, c) and 10 μm (c) confocal z-stacks are shown. Scale bars, 10 μm (a, lower panel b, c) and 25 μm (upper panel b, d).

**Extended data Figure 2. Imatinib has limited effect on immune cell infiltration after MCAO.**

(a) Immunofluorescence co-stainings of CD31 (endothelial cells, red) with MPO (neutrophils), CD3 (T cells) or B220 (B cells) (all in green), respectively at different time points after MCAO. Two headed arrows depict the respective immune cells transmigrated into the brain parenchyma and arrows immune cells in the blood vessel lumen, respectively. Cell nuclei were visualized by DAPI (blue). Representative maximum intensity projections of 12 μm z-stacks are shown. Scale bar, 10 μm. (b) Quantifications of MPO, CD3 and B220 expression was determined by manual counting of positive cells in the whole ischemic area in four different sections per animal. Symbols represent individual data points and bars the group mean ± S.E.M. Statistical significance $**p < 0.01$ relative to control (one-way ANOVA with Fisher's LSD test).

**Extended data Figure 3. Imatinib controls the structure of the PDGFRα scar 7 days after MCAO.**
Confocal images of immunofluorescent co-stainings of PDGFRα with GFAP 7 days post MCAO show that the PDGFRα$^+$ scar (demarcated with dashed lines) is localized on the ischemic core side of the GFAP$^+$ scar and there is no apparent co-localization of PDGFRα with GFAP in the glia scar. Further the PDGFRα$^+$ scar appears unstructured and enlarged in vehicle controls compared to in imatinib-treated mice. Representative images are shown. Scale bars, 50 µm.

**Extended data Figure 4. Imatinib reduces myofibroblast detachment from the vasculature.**
(a) Immunofluorescent co-staining for AMSA (red) with the endothelial cell marker CD31 (white) in naïve wild type brain sections (vibratome free float stain). In the healthy brain, ASMA expression is only seen in vSMC (arrow) with no parenchymal expression (asterisk). (b) Co-staining for PDGFRα (green) and the endothelial cell marker CD31 (white) 7 days post MCAO. PDGFRα$^+$ cells stayed more firmly attached to the vessels in imatinib-treated mice (arrow) compared to in vehicle controls (two headed arrows). (c) Co-staining for ASMA and CD31 7 days post MCAO. Pronounced parenchymal ASMA expression (two headed arrows) and ASMA$^+$ cells seemingly leaving the vessel wall (arrowheads) are induced by MCAO and reduced by imatinib treatment. ASMA$^+$ vSMC appeared unaffected (arrows). Cell nuclei were visualized by DAPI (blue). Representative maximum intensity projections of 21 µm (a), 10 µm (b) and 9 µm (c) confocal z-stacks are shown. Scale bars, 10 µm (a) and 25 µm (b, c).

**Extended data Figure 5. Imatinib reduces MCAO-induced ECM deposition in the fibrotic rim but not in the lesion core.** Immunofluorescent staining of fibronectin on brain sections from imatinib-treated and vehicle control mice 7 days post MCAO. (a) Imatinib treatment reduced Fibronectin$^+$ signal in the highly nucleated scar region compared to vehicle controls (demarcated by the two dashed lines). (b) Higher magnification of boxed areas in a. Representative epifluorescent images are shown. Scale bars, 250 µm (a) and 25 µm (b).

**Supplementary table 1.** Differentially expressed genes 3 hours post MCAO (down or upregulated in imatinib samples). This table includes a list of differentially expressed genes from vascular fragments of vehicle compared to imatinib-treated mice 3 hours post MCAO. Genes with *P*-value < 0.05 and a log2 (0.5) fold change are shown. For each gene the association with fibrosis, inflammation or metabolism is indicated.

**Supplementary table 2.** Differentially expressed genes 24 hours post MCAO (down or upregulated in imatinib samples). This table includes a list of differentially expressed genes from vascular fragments of vehicle compared to imatinib-treated mice 24 hours post MCAO. Genes with *P*-value < 0.05 and a log2 (0.5) fold change are shown. For each gene the association with fibrosis, inflammation or metabolism is indicated.

**Supplementary table 3.** Comparison of inflammation and fibrosis associated genes 3h post MCAO depicted in Figure 2d and Supplementary table 1 with the single cell RNAseq database of mouse brain vascular and vessel-associated cell types [33] reveals that a high percentage of fibrosis and inflammation associated differentially expressed genes in our datasets is expressed in the NVU, especially in astrocytes and fibroblast like cells.

**Supplementary table 4.** Comparison 3h post MCAO to human data. Transcripts that were significantly regulated in vascular fragments isolated from imatinib-treated mice compared to vehicle controls at 3 hours post MCAO were compared to microarray data from human perihematomal area (GSE24265). 49% of the common genes for both data were differentially expressed in both human and our data set. For each gene the association with fibrosis, inflammation or metabolism is indicated.

**Supplementary table 5.** Comparison 24h post MCAO to human data. Transcripts that were significantly regulated in vascular fragments isolated from imatinib-treated mice compared to vehicle controls at 24 hours post MCAO were compared to microarray data from human perihematomal area (GSE24265). 48% of the common genes for both data were differentially expressed in both human and our data set. For each gene the association with fibrosis, inflammation or metabolism is indicated.

**Zeitelhofer et al. Supplementary table 1. 3h post MCAO imatinib (down or upregulated in imatinib samples)**

| Symbol | Entrez Gene Name | Affymetrix II | Fold change | p value | Type | Fibrosis association | Inflammation association | Metabolism association |
|---|---|---|---|---|---|---|---|---|
| Abcc9 | ATP-binding cassette, sub-family C (CFTR/MRP), member 9 | 17472536 | -0.613 | 4.40E-02 | ion channel | yes | | |
| Ada | adenosine deaminase | 17394063 | 0.547 | 1.14E-02 | enzyme | yes | yes | yes |
| Aldoc | aldolase C, fructose-bisphosphate | 17253599 | -0.545 | 2.88E-02 | enzyme | yes | | yes |
| Alyref | Aly/REF export factor | 17273191 | -0.550 | 1.28E-02 | other | | | |
| Ankrd49 | ankyrin repeat domain 49 | 17524087 | 0.504 | 1.41E-02 | transcription regulator | | | |
| Arhgap8/P | Rho GTPase activating protein 8 | 17313751 | 0.532 | 2.98E-02 | other | | | |
| Arnt2 | aryl-hydrocarbon receptor nuclear translocator 2 | 17492985 | -0.815 | 3.78E-04 | transcription regulator | yes | | |
| Asl | argininosuccinate lyase | 17453179 | -0.956 | 1.56E-04 | enzyme | yes | | yes |
| Atp5g1 | ATP synthase, H+ transporting, mitochondrial Fo complex, subunit C1 (subunit 9) | 17268289 | -0.603 | 1.07E-02 | transporter | yes | | |
| Atxn7l1 | ataxin 7-like 1 | 17274940 | -0.597 | 2.35E-02 | other | | | |
| Awat2 | acyl-CoA wax alcohol acyltransferase 2 | 17543460 | -0.506 | 5.05E-03 | other | | | |
| Bcas1 | breast carcinoma amplified sequence 1 | 17394943 | -0.621 | 1.55E-02 | other | | | |
| Cacna1e | calcium channel, voltage-dependent, R type, alpha 1E subunit | 17228293 | 0.503 | 1.05E-02 | ion channel | | | |
| Cass4 | Cas scaffolding protein family member 4 | 17380119 | -0.698 | 2.48E-04 | other | | | |
| Ccl2 | chemokine (C-C motif) ligand 2 | 17254059 | -0.522 | 3.14E-02 | cytokine | yes | yes | |
| Ccnb1 | cyclin B1 | 17490038 | 0.654 | 9.55E-03 | kinase | yes | | |
| Ccne1 | cyclin E1 | 17489886 | -0.702 | 1.55E-05 | transcription regulator | yes | | |
| Cd248 | CD248 molecule, endosialin | 17356369 | -0.515 | 3.94E-02 | other | | | |
| Cd55 | CD55 molecule, decay accelerating factor for complement (Cromer blood group) | 17226622 | 0.814 | 2.84E-02 | other | yes | yes | |
| Cdh11 | cadherin 11, type 2, OB-cadherin (osteoblast) | 17512153 | -0.545 | 4.83E-02 | other | yes | yes | |
| Cenpk | centromere protein K | 17289584 | 0.589 | 8.92E-03 | other | yes | | |
| Cenpu | centromere protein U | 17501025 | -0.607 | 1.01E-02 | other | yes | | |
| Cldn6 | claudin 6 | 17334097 | 1.269 | 3.20E-07 | other | | | |
| Cntfr | ciliary neurotrophic factor receptor | 17424279 | -0.505 | 4.32E-02 | transmembrane receptor | | yes | |
| Cxcl10 | chemokine (C-X-C motif) ligand 10 | 17449718 | -0.588 | 4.77E-03 | cytokine | yes | yes | |
| Cxorf40a/( | chromosome X open reading frame 40A | 17535274 | 0.595 | 4.47E-02 | other | | | |
| Cyp2c54 ( | cytochrome P450, family 2, subfamily c, polypeptide 54 | 17364452 | 0.566 | 6.23E-03 | enzyme | | | |
| Dapk2 | death-associated protein kinase 2 | 17518680 | -0.538 | 1.16E-02 | kinase | yes | yes | |
| Doc2b | double C2-like domains, beta | 17266010 | 0.590 | 1.27E-02 | transporter | | | yes |
| Esd | esterase D | 17296971 | 0.572 | 2.07E-02 | enzyme | yes | | yes |
| Fabp5 | fatty acid binding protein 5 (psoriasis-associated) | 17548717 | -0.511 | 2.34E-02 | transporter | yes | | yes |
| Fabp7 | fatty acid binding protein 7, brain | 17233384 | -0.831 | 9.45E-03 | transporter | yes | | yes |
| Fam210a | family with sequence similarity 210, member A | 17355256 | -0.532 | 5.69E-03 | other | | | |
| Fbxo2 | F-box protein 2 | 17421588 | -0.560 | 8.38E-03 | enzyme | | | |
| Fbxw21 (I | F-box and WD-40 domain protein 21 | 17531315 | 0.533 | 1.89E-02 | other | | | |
| Fgr | FGR proto-oncogene, Src family tyrosine kinase | 17419483 | 0.632 | 3.38E-02 | kinase | | yes | |
| Fjx1 | four jointed box 1 | 17388725 | -0.544 | 1.33E-02 | other | | | |
| Fut11 | fucosyltransferase 11 (alpha (1,3) fucosyltransferase) | 17297495 | 1.605 | 2.00E-02 | enzyme | | | yes |
| Fxyd6 | FXYD domain containing ion transport regulator 6 | 17516777 | -0.672 | 4.71E-02 | ion channel | | | |
| Gabrq | gamma-aminobutyric acid (GABA) A receptor, theta | 17535420 | -0.513 | 1.91E-02 | ion channel | | | |
| Gbp2 | guanylate binding protein 2, interferon-inducible | 17403268 | -0.558 | 2.34E-02 | enzyme | yes | | |
| Gbp4 | guanylate binding protein 4 | 17403237 | -0.520 | 3.78E-02 | enzyme | | | |
| Gbp5 | guanylate binding protein 5 | 17403205 | -0.625 | 1.35E-02 | enzyme | | | |
| Gfpt2 | glutamine-fructose-6-phosphate transaminase 2 | 17249036 | -0.528 | 4.60E-03 | enzyme | yes | | yes |
| Gm15807/ | high-mobility group nucleosome binding domain 5 | 17328333 | 0.634 | 1.36E-02 | other | | | |
| Gnao1 | guanine nucleotide binding protein (G protein), alpha activating activity polypeptide O | 17503884 | -0.612 | 5.73E-04 | enzyme | yes | | |
| Gpx8 | glutathione peroxidase 8 (putative) | 17296281 | -0.542 | 2.53E-02 | enzyme | | | |
| Grhl1 | grainyhead-like transcription factor 1 | 17274513 | 0.520 | 6.52E-03 | transcription regulator | yes | | |
| Grk4 | G protein-coupled receptor kinase 4 | 17436659 | 0.568 | 2.74E-02 | kinase | | | |
| Gstm2 | glutathione S-transferase mu 2 (muscle) | 17409099 | 0.549 | 1.71E-02 | enzyme | yes | | yes |
| Hba1/Hba2 | hemoglobin, alpha 1 | 17248276 | 0.708 | 3.21E-02 | transporter | | | yes |
| Hibadh | 3-hydroxyisobutyrate dehydrogenase | 17466919 | -0.677 | 1.58E-02 | enzyme | yes | | |
| Hist1h4i | histone cluster 1, H4i | 17285815 | 0.917 | 2.94E-02 | other | | | |
| Hla-A | major histocompatibility complex, class I, A | 17337089 | -0.513 | 1.58E-02 | other | yes | yes | |
| Hyal3 | hyaluronoglucosaminidase 3 | 17521448 | 0.549 | 2.86E-02 | enzyme | | | |
| Idi1 | isopentenyl-diphosphate delta isomerase 1 | 17285056 | -0.579 | 1.43E-02 | enzyme | yes | | yes |
| Ifi16 | interferon, gamma-inducible protein 16 | 17230111 | 0.879 | 3.88E-02 | transcription regulator | yes | yes | |
| Igdcc4 | immunoglobulin superfamily, DCC subclass, member 4 | 17518458 | -0.515 | 1.98E-03 | other | | yes | |
| Igtp | interferon gamma induced GTPase | 17249980 | -0.714 | 1.59E-02 | enzyme | | yes | |
| Irs1 | insulin receptor substrate 1 | 17214753 | -0.516 | 1.60E-02 | enzyme | yes | | |
| Itpk1 | inositol-tetrakisphosphate 1-kinase | 17283463 | -0.660 | 3.75E-03 | kinase | | | |
| Kcnk2 | potassium channel, two pore domain subfamily K, member 2 | 17230902 | 0.518 | 7.34E-03 | ion channel | | | |
| Kiaa0930 | KIAA0930 | 17320035 | -0.825 | 8.13E-03 | other | | | |
| Krt27 | keratin 27, type I | 17269042 | 0.528 | 4.69E-03 | other | | | |
| Lhfp | lipoma HMGIC fusion partner | 17397511 | -0.510 | 1.44E-02 | other | | | |
| Ly6a (Incl | lymphocyte antigen 6 complex, locus A | 17312219 | -0.506 | 1.46E-02 | other | | | |
| Mbp | myelin basic protein | 17352175 | -0.957 | 9.28E-03 | other | yes | | |
| Mir-194 | microRNA 194-1 | 17356739 | -0.524 | 8.69E-03 | microRNA | | | |
| Mir-290 | microRNA 372 | 17473111 | 1.226 | 1.46E-02 | microRNA | | | |
| Mir-432 | microRNA 432 | 17278711 | 0.861 | 1.68E-04 | microRNA | | | |
| Mir-467 | microRNA 466 | 17366718 | -0.554 | 2.08E-03 | microRNA | | | |
| Mir-500 | microRNA 501 | 17539820 | 0.517 | 7.22E-03 | microRNA | | | |
| Mir-743 | microRNA 890 | 17542062 | -0.594 | 1.20E-02 | microRNA | | | |
| Mob3a | MOB kinase activator 3A | 17235396 | -0.527 | 3.57E-04 | other | | | |
| Mobp | myelin-associated oligodendrocyte basic protein | 17523192 | -0.727 | 2.39E-03 | other | | | |
| Ms4a4b (I | membrane-spanning 4-domains, subfamily A, member 4B | 17357648 | -0.809 | 4.42E-02 | other | | yes | |
| Mt3 | metallothionein 3 | 17503932 | -0.646 | 3.20E-02 | other | yes | | |
| Myoc | myocilin, trabecular meshwork inducible glucocorticoid response | 17218752 | -0.633 | 3.35E-02 | other | | | |
| Nanp | N-acetylneuraminic acid phosphatase | 17340397 | -1.390 | 9.99E-03 | enzyme | | | |
| Nkain2 | Na+/K+ transporting ATPase interacting 2 | 17240109 | -0.576 | 3.41E-02 | other | | | |
| Nmrk1 | nicotinamide riboside kinase 1 | 17358020 | 0.600 | 1.48E-02 | kinase | | | |
| Nov | nephroblastoma overexpressed | 17311519 | 0.602 | 1.75E-03 | growth factor | | yes | |
| Npr3 | natriuretic peptide receptor 3 | 17316043 | 0.589 | 1.55E-02 | G-protein coupled recepto | yes | | |
| Nsun6 | NOP2/Sun domain family, member 6 | 17381911 | 0.512 | 3.84E-02 | enzyme | | | |
| Ott (Includ | ovary testis transcribed | 17538452 | -0.671 | 2.16E-02 | other | | | |
| Pdgfra | platelet-derived growth factor receptor, alpha polypeptide | 17438246 | -0.828 | 1.02E-03 | kinase | yes | yes | |
| Poldip2 | polymerase (DNA-directed), delta interacting protein 2 | 17253674 | -0.520 | 5.18E-04 | other | | | |
| Pou2f3 | POU class 2 homeobox 3 | 17526086 | 0.583 | 6.76E-04 | transcription regulator | | | |
| Ppargc1a | peroxisome proliferator-activated receptor gamma, coactivator 1 alpha | 17448001 | -0.593 | 6.21E-03 | transcription regulator | yes | | yes |
| Ppp1r14a | protein phosphatase 1, regulatory (inhibitor) subunit 14A | 17476119 | -0.683 | 2.90E-02 | phosphatase | | | |
| Psmb9 | proteasome subunit beta 9 | 17343789 | -0.521 | 2.81E-02 | peptidase | yes | | |
| Rdh16 | retinol dehydrogenase 16 (all-trans) | 17238172 | -0.559 | 7.43E-03 | enzyme | yes | | |
| Rdh5 | retinol dehydrogenase 5 (11-cis/9-cis) | 17246358 | 0.506 | 4.22E-02 | enzyme | | | |
| Rell2 | RELT-like 2 | 17349962 | 0.549 | 2.58E-02 | other | | | |
| Rhbdl2 | rhomboid, veinlet-like 2 (Drosophila) | 17418223 | -0.558 | 2.70E-02 | peptidase | | | |
| Rnf113a1 | ring finger protein 113A1 | 17534157 | 0.543 | 3.12E-02 | other | | yes | |
| Rpp30 | ribonuclease P/MRP 30kDa subunit | 17358876 | 0.566 | 4.64E-03 | enzyme | | | |
| Scn10a | sodium channel, voltage gated, type X alpha subunit | 17532184 | -0.533 | 3.67E-03 | ion channel | | | |
| Serf1a/Serf | small EDRK-rich factor 1A (telomeric) | 17289455 | -0.522 | 1.90E-03 | other | | | |
| Shisa9 | shisa family member 9 | 17322985 | -0.536 | 1.06E-02 | other | | | |
| Sirpb1 | signal-regulatory protein beta 1 | 17404200 | -0.553 | 4.33E-02 | other | | yes | |
| Slc16a12 | solute carrier family 16, member 12 | 17364139 | -0.527 | 3.98E-02 | transporter | | | |
| Slc9a2 | solute carrier family 9, subfamily A (NHE2, cation proton antiporter 2), member 2 | 17212286 | 0.592 | 2.20E-02 | transporter | | | |
| Slc9a7 | solute carrier family 9, subfamily A (NHE7, cation proton antiporter 7), member 7 | 17540465 | -0.545 | 3.71E-02 | transporter | | | |
| Slco1a2 | solute carrier organic anion transporter family, member 1A2 | 17472443 | -0.561 | 2.56E-02 | transporter | | | |
| Snhg11 | small nucleolar RNA host gene 11 | 17378848 | -0.639 | 2.63E-03 | other | | | |
| Strip2 | striatin interacting protein 2 | 17456772 | 0.868 | 2.84E-03 | other | | | |
| Tatdn3 | TatD DNase domain containing 3 | 17231015 | -0.521 | 6.08E-03 | other | | | |
| Tfap2b | transcription factor AP-2 beta (activating enhancer binding protein 2 beta) | 17211347 | 0.609 | 5.72E-03 | transcription regulator | | | |
| Tfb2m | transcription factor B2, mitochondrial | 17230331 | 0.932 | 2.53E-03 | enzyme | yes | | |
| Thumpd2 | THUMP domain containing 2 | 17347606 | 0.732 | 1.09E-02 | enzyme | | | |
| Tmem181 | transmembrane protein 181 | 17332893 | -0.529 | 7.36E-04 | other | | | |
| Tmsb4x (I | thymosin, beta 4, X chromosome | 17544786 | -0.639 | 1.68E-02 | other | yes | | |
| Tnf | tumor necrosis factor | 17344309 | 0.588 | 4.18E-02 | cytokine | yes | yes | |
| Trabd2b | TraB domain containing 2B | 17417090 | -0.645 | 1.49E-02 | peptidase | | | |
| Uchl1 | ubiquitin carboxyl-terminal esterase L1 (ubiquitin thiolesterase) | 17437876 | -0.523 | 6.41E-04 | peptidase | yes | | |
| Vmn1r180 | vomeronasal 1 receptor 180 | 17487533 | -0.784 | 5.39E-03 | other | | | |
| Zbtb45 | zinc finger and BTB domain containing 45 | 17486603 | 0.500 | 8.12E-04 | other | | | |
| Znf25 | zinc finger protein 25 | 17470187 | 0.508 | 3.98E-03 | other | | | |
| Znf446 | zinc finger protein 446 | 17473817 | -0.508 | 3.84E-04 | transcription regulator | | | |
| Znf676 | zinc finger protein 676 | 17288344 | 0.585 | 1.95E-02 | other | | | |

**Zeitelhofer et al supplemental table 2. 24h post MCAO (down or upregulated in imatinib samples)**

| Symbol | Entrez Gene Name | Affymetrix ID | Fold change | p value | Type | Fibrosis association | Inflammation association | Metabolism association |
|---|---|---|---|---|---|---|---|---|
| Abhd18 | abhydrolase domain containing 18 | 17397297 | 0.580 | 2.33E-02 | other | | | |
| Adcy3 | adenylate cyclase 3 | 17273714 | 0.547 | 3.95E-03 | enzyme | | | yes |
| Aldob | aldolase B, fructose-bisphosphate | 17425233 | -0.738 | 3.63E-04 | enzyme | yes | | yes |
| Atp2b3 | ATPase, Ca++ transporting, plasma membrane 3 | 17535572 | 0.692 | 4.15E-02 | transporter | | | |
| C3 | complement component 3 | 17346528 | -0.634 | 4.25E-02 | peptidase | yes | yes | |
| Ca6 | carbonic anhydrase VI | 17433265 | -0.628 | 8.17E-05 | enzyme | | | yes |
| Ca8 | carbonic anhydrase VIII | 17423121 | 0.505 | 4.18E-02 | enzyme | | | yes |
| Ccbl1 | cysteine conjugate-beta lyase, cytoplasmic | 17383588 | 0.562 | 6.41E-05 | enzyme | yes | | yes |
| Ccl22 | chemokine (C-C motif) ligand 22 | 17504122 | -0.878 | 5.02E-03 | cytokine | yes | yes | |
| Ccl5 | chemokine (C-C motif) ligand 5 | 17266946 | -0.632 | 1.86E-02 | cytokine | yes | yes | |
| Cd200r1 | CD200 receptor 1 | 17325861 | -0.513 | 1.06E-02 | other | | | |
| Cenpq | centromere protein Q | 17344990 | 0.549 | 2.02E-02 | other | | | |
| Colgalt2 | collagen beta(1-O)galactosyltransferase 2 | 17218233 | 0.534 | 1.14E-02 | other | | | |
| Cox6a2 | cytochrome c oxidase subunit VIa polypeptide 2 | 17496857 | -0.547 | 1.92E-02 | enzyme | yes | | |
| Cshl1 | chorionic somatomammotropin hormone-like 1 | 17270829 | 1.769 | 3.80E-02 | transcription regulator | | | |
| Dsg1 | desmoglein 1 | 17348726 | -0.530 | 1.10E-02 | other | | | |
| Egln3 | egl-9 family hypoxia-inducible factor 3 | 17281084 | -0.850 | 2.07E-02 | enzyme | yes | | |
| Eps8l2 | EPS8-like 2 | 17484852 | 0.672 | 7.57E-05 | other | yes | | |
| F10 | coagulation factor X | 17499224 | -0.596 | 2.35E-02 | peptidase | | | |
| Fgr | FGR proto-oncogene, Src family tyrosine kinase | 17419483 | -0.733 | 8.49E-03 | kinase | | yes | |
| Fhit | fragile histidine triad | 17303474 | -0.563 | 8.36E-05 | enzyme | yes | | |
| Gabarapl1 | GABA(A) receptor-associated protein like 1 | 17463550 | 0.614 | 4.04E-04 | other | yes | | |
| Gpat3 | glycerol-3-phosphate acyltransferase 3 | 17439622 | -0.504 | 4.92E-03 | enzyme | | | yes |
| Gpc3 | glypican 3 | 17541681 | 0.661 | 9.08E-03 | other | yes | | |
| Hba1/Hba2 | hemoglobin, alpha 1 | 17248276 | -0.633 | 3.21E-02 | transporter | | | |
| Hist1h4c | histone cluster 1, H4c | 17408021 | 0.593 | 3.98E-02 | other | | | |
| Hla-Dqa1 | major histocompatibility complex, class II, DQ alpha 1 | 17343813 | -0.826 | 3.33E-02 | transmembrane receptor | yes | yes | |
| Hla-Drb5 | major histocompatibility complex, class II, DR beta 5 | 17336502 | -0.846 | 2.39E-02 | transmembrane receptor | | yes | |
| Hmga1 | high mobility group AT-hook 1 | 17335129 | 0.592 | 1.23E-02 | transcription regulator | | | |
| Hpse | heparanase | 17548411 | -1.058 | 1.82E-02 | enzyme | yes | | |
| Ifi16 | interferon, gamma-inducible protein 16 | 17230067 | -0.514 | 2.05E-02 | transcription regulator | yes | yes | |
| Il1r2 | interleukin 1 receptor, type II | 17212174 | -0.693 | 4.23E-02 | transmembrane receptor | yes | yes | |
| Il1rap | interleukin 1 receptor accessory protein | 17324542 | -0.547 | 8.09E-04 | transmembrane receptor | yes | yes | |
| Itgax | integrin, alpha X (complement component 3 receptor 4 subunit) | 17483615 | -0.878 | 7.90E-03 | transmembrane receptor | yes | yes | |
| Kcnn4 | potassium channel, calcium activated intermediate/small conductance subfamily N alpha, member 4 | 17474941 | -0.568 | 1.06E-02 | ion channel | | | |
| Kiaa0930 | KIAA0930 | 17320035 | 0.546 | 3.96E-02 | other | | | |
| Kif23 | kinesin family member 23 | 17527934 | 0.514 | 9.69E-03 | other | yes | | |
| Klk3 | kallikrein-related peptidase 3 | 17477347 | -0.611 | 9.23E-03 | peptidase | | | |
| Klrd1 | killer cell lectin-like receptor subfamily D, member 1 | 17463567 | -0.949 | 1.08E-03 | transmembrane receptor | | | |
| Klrk1 | killer cell lectin-like receptor subfamily K, member 1 | 17471565 | -0.778 | 9.49E-04 | other | | | |
| Kmo | kynurenine 3-monooxygenase (kynurenine 3-hydroxylase) | 17219789 | -0.626 | 1.20E-02 | enzyme | | | |
| Lepr | leptin receptor | 17415979 | 0.541 | 9.00E-03 | transmembrane receptor | yes | | yes |
| Lilrb3 | leukocyte immunoglobulin-like receptor, subfamily B (with TM and ITIM domains), member 3 | 17485589 | -0.684 | 2.29E-02 | transmembrane receptor | yes | yes | |
| Ltb4r | leukotriene B4 receptor | 17300666 | -0.606 | 1.83E-02 | G-protein coupled receptor | | yes | |
| Mbp | myelin basic protein | 17352175 | 0.643 | 4.06E-02 | other | yes | | |
| Mia | melanoma inhibitory activity | 17488151 | 0.638 | 1.51E-02 | other | | | |
| Mir-142 | microRNA 142 | 17254838 | -0.894 | 6.53E-03 | microRNA | | | |
| Mir-15 | microRNA 15a | 17307518 | -0.684 | 4.96E-02 | microRNA | | | |
| Mir-150 | microRNA 150 | 17477693 | 0.589 | 2.88E-02 | microRNA | | | |
| Mir-181 | microRNA 181a-1 | 17217846 | 0.542 | 6.30E-03 | microRNA | | | |
| Mir-27 | microRNA 27a | 17503118 | -0.595 | 2.63E-02 | microRNA | | | |
| Ms4a4b (Includ | membrane-spanning 4-domains, subfamily A, member 4B | 17357648 | -0.835 | 2.29E-02 | other | | yes | |
| Napsa | napsin A aspartic peptidase | 17477508 | -0.748 | 1.19E-02 | peptidase | | | |
| Nid1 | nidogen 1 | 17285225 | 0.764 | 2.24E-03 | other | yes | | |
| Otx2 | orthodenticle homeobox 2 | 17305830 | 0.544 | 4.27E-02 | transcription regulator | | | |
| Palm | paralemmin | 17234963 | 0.650 | 1.17E-04 | other | | | |
| Pcolce2 | procollagen C-endopeptidase enhancer 2 | 17520425 | 0.605 | 2.13E-02 | other | | | |
| Pdcd1lg2 | programmed cell death 1 ligand 2 | 17358552 | -0.596 | 3.70E-03 | enzyme | | | |
| Pde2a | phosphodiesterase 2A, cGMP-stimulated | 17480880 | 0.567 | 1.93E-04 | enzyme | | | yes |
| Pdgfd | platelet derived growth factor D | 17514460 | 0.509 | 7.26E-03 | growth factor | yes | | |
| Peg3 | paternally expressed 3 | 17486110 | 0.602 | 1.09E-02 | kinase | | | |
| Pilra | paired immunoglobin-like type 2 receptor alpha | 17454166 | -0.703 | 1.92E-02 | other | | | |
| Plppr4 | phospholipid phosphatase related 4 | 17409816 | 0.559 | 2.49E-02 | phosphatase | | | yes |
| Poc5 | POC5 centriolar protein | 17289289 | 0.537 | 7.00E-03 | other | | | |
| Prl | prolactin | 17286107 | 1.717 | 4.51E-02 | cytokine | yes | | |
| Prrg1 | proline rich Gla (G-carboxyglutamic acid) 1 | 17542948 | 0.568 | 9.91E-05 | other | | | yes |
| Rab20 | RAB20, member RAS oncogene family | 17507435 | -0.508 | 4.75E-03 | enzyme | yes | | |
| Rab7b | RAB7B, member RAS oncogene family | 17217048 | 0.555 | 4.91E-03 | peptidase | | | |
| Rasl11b | RAS-like, family 11, member B | 17438189 | 0.675 | 1.74E-02 | enzyme | yes | | |
| Rhbdl2 | rhomboid, veinlet-like 2 (Drosophila) | 17418223 | 0.635 | 7.04E-03 | peptidase | | | |
| Runx2 | runt-related transcription factor 2 | 17345134 | -0.834 | 2.67E-03 | transcription regulator | yes | | |
| Serpinb1b | serine (or cysteine) peptidase inhibitor, clade B, member 1b | 17286385 | -0.528 | 4.78E-03 | enzyme | yes | | |
| Serpinh1 | serpin peptidase inhibitor, clade H (heat shock protein 47), member 1, (collagen binding protein 1) | 17493658 | 0.581 | 8.47E-04 | other | yes | yes | |
| Sgtb | small glutamine-rich tetratricopeptide repeat (TPR)-containing, beta | 17289553 | 0.534 | 1.51E-03 | other | | | |
| Sh3bgrl | SH3 domain binding glutamate-rich protein like | 17537199 | -0.828 | 7.31E-03 | other | | | |
| Sirpb1 | signal-regulatory protein beta 1 | 17404217 | -1.521 | 2.15E-06 | other | | yes | |
| Slamf7 | SLAM family member 7 | 17229782 | -0.819 | 4.47E-02 | other | | | |
| Slc4a10 | solute carrier family 4, sodium bicarbonate transporter, member 10 | 17371162 | 0.694 | 1.88E-02 | transporter | | | |
| Slc4a5 | solute carrier family 4 (sodium bicarbonate cotransporter), member 5 | 17460023 | 0.530 | 2.76E-02 | transporter | | | |
| Slfn12l | schlafen family member 12-like | 17254176 | -0.737 | 3.76E-02 | enzyme | | | |
| Spry2 | sprouty RTK signaling antagonist 2 | 17309340 | 0.530 | 1.90E-02 | other | yes | | |
| Strip2 | striatin interacting protein 2 | 17456772 | 0.525 | 2.93E-02 | other | | | |
| Taf1d | TATA box binding protein (TBP)-associated factor, RNA polymerase I, D, 41kDa | 17514834 | -0.666 | 4.84E-02 | other | yes | | |
| Trem3 | triggering receptor expressed on myeloid cells 3 | 17338371 | -0.623 | 1.03E-03 | other | | | |
| Vwa1 | von Willebrand factor A domain containing 1 | 17433879 | 0.531 | 4.08E-05 | other | | | |

**Zeitelhofer et al Supplemental table 3. Cellular expression of the fibrosis- and inflammation associated genes regulated 3h post MCAO by imatinib according to the single cell RNAseq database of mouse brain vascular and vessel-associated cell types (http://betsholtzlab.org/VascularSingleCells/database.html)**

| Symbol | Entrez Gene Name | Affymetrix ID | Fd change | p value | Type | Fibrosis association | Inflammation association | NVU cell type specific expression |
|---|---|---|---|---|---|---|---|---|
| Abcc9 | ATP-binding cassette, sub-family C (CFTR/MRP), member 9 | 17472536 | -0.613 | 4.40E-02 | ion channel | yes | | FB2, PC, SMC |
| Ada | adenosine deaminase | 17394063 | 0.547 | 1.14E-02 | enzyme | yes | yes | EC |
| Aldoc | aldase C, fructose-bisphosphate | 17253599 | -0.545 | 2.88E-02 | enzyme | yes | | AC |
| Arnt2 | aryl-hydrocarbon receptor nuclear translocator 2 | 17492985 | -0.815 | 3.78E-04 | transcription regulator | yes | | AC, FB1/2, PC |
| Asl | argininosuccinate lyase | 17453179 | -0.956 | 1.56E-04 | enzyme | yes | | All |
| Atp5g1 | ATP synthase, H+ transporting, mitochondrial Fo complex, subunit C1 (subunit 9) | 17268289 | -0.603 | 1.07E-02 | transporter | yes | | All |
| Ccl2 | chemokine (C-C motif) ligand 2 | 17254059 | -0.522 | 3.14E-02 | cytokine | yes | yes | Not in unchallenged NVU |
| Ccnb1 | cyclin B1 | 17490038 | 0.654 | 9.55E-03 | kinase | yes | | Not detected in the dataset |
| Ccne1 | cyclin E1 | 17489886 | -0.702 | 1.55E-05 | transcription regulator | yes | | Not in unchallenged NVU |
| Cd55 | CD55 mecule, decay accelerating factor for complement (Cromer blood group) | 17226622 | 0.814 | 2.84E-02 | other | yes | yes | FB1, EC |
| Cdh11 | cadherin 11, type 2, OB-cadherin (osteoblast) | 17512153 | -0.545 | 4.83E-02 | other | yes | yes | AC, FB1/2, PC, SMC |
| Cenpk | centromere protein K | 17289584 | 0.589 | 8.92E-03 | other | yes | | Not in unchallenged NVU |
| Cenpu | centromere protein U | 17501025 | -0.607 | 1.01E-02 | other | yes | | Not in unchallenged NVU |
| Cntfr | ciliary neurotrophic factor receptor | 17424279 | -0.505 | 4.32E-02 | transmembrane receptor | | yes | AC, FB1/2 |
| Cxcl10 | chemokine (C-X-C motif) ligand 10 | 17449718 | -0.588 | 4.77E-03 | cytokine | yes | yes | FB2 |
| Dapk2 | death-associated protein kinase 2 | 17518680 | -0.538 | 1.16E-02 | kinase | yes | yes | FB1/2 |
| Esd | esterase D | 17296971 | 0.572 | 2.07E-02 | enzyme | yes | | All |
| Fabp5 | fatty acid binding protein 5 (psoriasis-associated) | 17548717 | -0.511 | 2.34E-02 | transporter | yes | | AC, PC, FB2, SMC |
| Fabp7 | fatty acid binding protein 7, brain | 17233384 | -0.831 | 9.45E-03 | transporter | yes | | AC |
| Fgr | FGR proto-oncogene, Src family tyrosine kinase | 17419483 | 0.632 | 3.38E-02 | kinase | | yes | Not in unchallenged NVU |
| Gbp2 | guanylate binding protein 2, interferon-inducible | 17403268 | -0.558 | 2.34E-02 | enzyme | yes | | Not in unchallenged NVU |
| Gfpt2 | glutamine-fructose-6-phosphate transaminase 2 | 17249036 | -0.528 | 4.60E-03 | enzyme | yes | | AC, PC, FB1/2 |
| Gnao1 | guanine nucleotide binding protein (G protein), alpha activating activity pypeptide O | 17503884 | -0.612 | 5.73E-04 | enzyme | yes | | AC, SMC, FB1/2, PC |
| Grhl1 | grainyhead-like transcription factor 1 | 17274513 | 0.520 | 6.52E-03 | transcription regulator | yes | | AC, EC |
| Gstm2 | glutathione S-transferase mu 2 (muscle) | 17409099 | 0.549 | 1.71E-02 | enzyme | yes | | EC, PC, FB1/2, SMC |
| Hibadh | 3-hydroxyisobutyrate dehydrogenase | 17466919 | -0.677 | 1.58E-02 | enzyme | yes | | All |
| Hla-A | major histocompatibility complex, class I, A | 17337089 | -0.513 | 1.58E-02 | other | yes | yes | Not detected in the dataset |
| Idi1 | isopentenyl-diphosphate delta isomerase 1 | 17285056 | -0.579 | 1.43E-02 | enzyme | yes | | AC, FB1 |
| Ifi16 | interferon, gamma-inducible protein 16 | 17230111 | 0.879 | 3.88E-02 | transcription regulator | yes | yes | Not in unchallenged NVU |
| Igdcc4 | immunoglobulin superfamily, DCC subclass, member 4 | 17518458 | -0.515 | 1.98E-03 | other | | yes | AC, FB1/2 |
| Igtp | interferon gamma induced GTPase | 17249980 | -0.714 | 1.59E-02 | enzyme | | yes | AC, EC, FB2 |
| Irs1 | insulin receptor substrate 1 | 17214753 | -0.516 | 1.60E-02 | enzyme | yes | | AC, PC, SMC, FB1/2 |
| Mbp | myelin basic protein | 17352175 | -0.957 | 9.28E-03 | other | yes | | AC |
| Ms4a4b (Incl | membrane-spanning 4-domains, subfamily A, member 4B | 17357648 | -0.809 | 4.42E-02 | other | | yes | Not in unchallenged NVU |
| Mt3 | metallothionein 3 | 17503932 | -0.646 | 3.20E-02 | other | yes | | AC |
| Nov | nephroblastoma overexpressed | 17311519 | 0.602 | 1.75E-03 | growth factor | | yes | Not in unchallenged NVU |
| Npr3 | natriuretic peptide receptor 3 | 17316043 | 0.589 | 1.55E-02 | G-protein coupled receptor | yes | | FB1/2 |
| Pdgfra | platelet-derived growth factor receptor, alpha pypeptide | 17438246 | -0.828 | 1.02E-03 | kinase | yes | yes | FB1/2 |
| Ppargc1a | peroxisome priferator-activated receptor gamma, coactivator 1 alpha | 17448001 | -0.593 | 6.21E-03 | transcription regulator | yes | | AC |
| Psmb9 | proteasome subunit beta 9 | 17343789 | -0.521 | 2.81E-02 | peptidase | yes | | EC, PC, FB1/2, SMC |
| Rdh16 | retin dehydrogenase 16 (all-trans) | 17238172 | -0.559 | 7.43E-03 | enzyme | yes | | Not in unchallenged NVU |
| Rnf113a1 | ring finger protein 113A1 | 17534157 | 0.543 | 3.12E-02 | other | | yes | Not in unchallenged NVU |
| Sirpb1 | signal-regulatory protein beta 1 | 17404200 | -0.553 | 4.33E-02 | other | | yes | Not in unchallenged NVU |
| Tfb2m | transcription factor B2, mitochondrial | 17230331 | 0.932 | 2.53E-03 | enzyme | yes | | All |
| Tmsb4x (Incl | thymosin, beta 4, X chromosome | 17544786 | -0.639 | 1.68E-02 | other | yes | | All |
| Tnf | tumor necrosis factor | 17344309 | 0.588 | 4.18E-02 | cytokine | yes | yes | Not in unchallenged NVU |
| Uchl1 | ubiquitin carboxyl-terminal esterase L1 (ubiquitin thiesterase) | 17437876 | -0.523 | 6.41E-04 | peptidase | yes | | PC, AC, SMC, EC |

Abbreviations: PC - Pericytes; SMC - Smooth muscle cells; FB - Vascular fibroblast-like cells;  - igodendrocytes; EC - Endothelial cells; AC - Astrocytes

**Zeitelhofer et al Supplemental table 4. 3h post MCAO imatinib (down or upregulated in imatinib samples) compared to microarray data from stroke patients (GSE24265)**

**Anticorrelation between datasets**

| Symbol | Entrez Gene Name | Affymetrix ID | Fold change | p value | Type | Fibrosis association | Inflammation association | Metabolism association |
|---|---|---|---|---|---|---|---|---|
| Ankrd49 | ankyrin repeat domain 49 | 17524087 | 0.504 | 1.41E-02 | transcription regulator | | | |
| Asl | argininosuccinate lyase | 17453179 | -0.956 | 1.56E-04 | enzyme | yes | | yes |
| Bcas1 | breast carcinoma amplified sequence 1 | 17394943 | -0.621 | 1.55E-02 | other | | | |
| Cacna1e | calcium channel, voltage-dependent, R type, alpha 1E subunit | 17228293 | 0.503 | 1.05E-02 | ion channel | | | |
| Ccl2 | chemokine (C-C motif) ligand 2 | 17254059 | -0.522 | 3.14E-02 | cytokine | yes | yes | |
| Ccnb1 | cyclin B1 | 17490038 | 0.654 | 9.55E-03 | kinase | yes | | |
| Fut11 | fucosyltransferase 11 (alpha (1,3) fucosyltransferase) | 17297495 | 1.605 | 2.00E-02 | enzyme | | | yes |
| Grhl1 | grainyhead-like transcription factor 1 | 17274513 | 0.520 | 6.52E-03 | transcription regulator | yes | | |
| Gstm2 | glutathione S-transferase mu 2 (muscle) | 17409099 | 0.549 | 1.71E-02 | enzyme | yes | | yes |
| Hibadh | 3-hydroxyisobutyrate dehydrogenase | 17466919 | -0.677 | 1.58E-02 | enzyme | yes | | |
| Hist1h4i | histone cluster 1, H4i | 17285815 | 0.917 | 2.94E-02 | other | | | |
| Hla-A | major histocompatibility complex, class I, A | 17337089 | -0.513 | 1.58E-02 | other | yes | yes | |
| Mobp | myelin-associated oligodendrocyte basic protein | 17523192 | -0.727 | 2.39E-03 | other | | | |
| Nmrk1 | nicotinamide riboside kinase 1 | 17358020 | 0.600 | 1.48E-02 | kinase | | | |
| Nov | nephroblastoma overexpressed | 17311519 | 0.602 | 1.75E-03 | growth factor | | yes | |
| Nsun6 | NOP2/Sun domain family, member 6 | 17381911 | 0.512 | 3.84E-02 | enzyme | | | |
| Poldip2 | polymerase (DNA-directed), delta interacting protein 2 | 17253674 | -0.520 | 5.18E-04 | other | | | |
| Ppp1r14a | protein phosphatase 1, regulatory (inhibitor) subunit 14A | 17476119 | -0.683 | 2.90E-02 | phosphatase | | | |
| Thumpd2 | THUMP domain containing 2 | 17347606 | 0.732 | 1.09E-02 | enzyme | | | |
| Tmsb4x (Incl | thymosin, beta 4, X chromosome | 17544786 | -0.639 | 1.68E-02 | other | yes | | |
| Znf25 | zinc finger protein 25 | 17470187 | 0.508 | 3.98E-03 | other | | | |

**Correlation between datasets**

| Symbol | Entrez Gene Name | Affymetrix ID | Fold change | p value | Type | Fibrosis association | Inflammation association | Metabolism association |
|---|---|---|---|---|---|---|---|---|
| Abcc9 | ATP-binding cassette, sub-family C (CFTR/MRP), member 9 | 17472536 | -0.613 | 4.40E-02 | ion channel | yes | | |
| Aldoc | aldolase C, fructose-bisphosphate | 17253599 | -0.545 | 2.88E-02 | enzyme | yes | | yes |
| Arnt2 | aryl-hydrocarbon receptor nuclear translocator 2 | 17492985 | -0.815 | 3.78E-04 | transcription regulator | yes | | |
| Atp5g1 | ATP synthase, H+ transporting, mitochondrial Fo complex, subunit C1 (subunit 9) | 17268289 | -0.603 | 1.07E-02 | transporter | yes | | |
| Cass4 | Cas scaffolding protein family member 4 | 17380119 | -0.698 | 2.48E-04 | other | | | |
| Cd55 | CD55 molecule, decay accelerating factor for complement (Cromer blood group) | 17226622 | 0.814 | 2.84E-02 | other | yes | yes | |
| Cdh11 | cadherin 11, type 2, OB-cadherin (osteoblast) | 17512153 | -0.545 | 4.83E-02 | other | yes | yes | |
| Fabp7 | fatty acid binding protein 7, brain | 17233384 | -0.831 | 9.45E-03 | transporter | yes | | yes |
| Fxyd6 | FXYD domain containing ion transport regulator 6 | 17516777 | -0.672 | 4.71E-02 | ion channel | | | |
| Hba1/Hba2 | hemoglobin, alpha 1 | 17248276 | 0.708 | 3.21E-02 | transporter | | | yes |
| Nanp | N-acetylneuraminic acid phosphatase | 17340397 | -1.390 | 9.99E-03 | enzyme | | | |
| Pdgfra | platelet-derived growth factor receptor, alpha polypeptide | 17438246 | -0.828 | 1.02E-03 | kinase | yes | yes | |
| Ppargc1a | peroxisome proliferator-activated receptor gamma, coactivator 1 alpha | 17448001 | -0.593 | 6.21E-03 | transcription regulator | yes | | yes |
| Tatdn3 | TatD DNase domain containing 3 | 17231015 | -0.521 | 6.08E-03 | other | | | |

**Zeitelhofer et al Supplemental table 5. 24h post MCAO (down or upregulated in imatinib samples) compared to microarray data from stroke patients (GSE24265)**

**Anticorrelation between datasets**

| Symbol | Entrez Gene Name | Affymetrix ID | Fold change | p value | Type | Fibrosis association | Inflammation association | Metabolism association |
|---|---|---|---|---|---|---|---|---|
| Adcy3 | adenylate cyclase 3 | 17273714 | 0.547 | 3.95E-03 | enzyme | | | yes |
| Ccl5 | chemokine (C-C motif) ligand 5 | 17266946 | -0.632 | 1.86E-02 | cytokine | yes | yes | |
| Colgalt2 | collagen beta(1-O)galactosyltransferase 2 | 17218233 | 0.534 | 1.14E-02 | other | | | |
| Fgr | FGR proto-oncogene, Src family tyrosine kinase | 17419483 | -0.733 | 8.49E-03 | kinase | | yes | |
| Hba1/Hba2 | hemoglobin, alpha 1 | 17248276 | -0.633 | 3.21E-02 | transporter | | | |
| Hla-Dqa1 | major histocompatibility complex, class II, DQ alpha 1 | 17343813 | -0.826 | 3.33E-02 | transmembrane receptor | yes | yes | |
| Hla-Drb5 | major histocompatibility complex, class II, DR beta 5 | 17336502 | -0.846 | 2.39E-02 | transmembrane receptor | | yes | |
| Il1r2 | interleukin 1 receptor, type II | 17212174 | -0.693 | 4.23E-02 | transmembrane receptor | yes | yes | |
| Itgax | integrin, alpha X (complement component 3 receptor 4 subunit) | 17483615 | -0.878 | 7.90E-03 | transmembrane receptor | yes | yes | |
| Lilrb3 | leukocyte immunoglobulin-like receptor, subfamily B (with TM and ITIM domains), member | 17485589 | -0.684 | 2.29E-02 | transmembrane receptor | yes | yes | |
| Mbp | myelin basic protein | 17352175 | 0.643 | 4.06E-02 | other | yes | | |
| Palm | paralemmin | 17234963 | 0.650 | 1.17E-04 | other | | | |
| Pde2a | phosphodiesterase 2A, cGMP-stimulated | 17480880 | 0.567 | 1.93E-04 | enzyme | | | yes |
| Pdgfd | platelet derived growth factor D | 17514460 | 0.509 | 7.26E-03 | growth factor | yes | | |
| Peg3 | paternally expressed 3 | 17486110 | 0.602 | 1.09E-02 | kinase | | | |
| Rasl11b | RAS-like, family 11, member B | 17438189 | 0.675 | 1.74E-02 | enzyme | yes | | |
| Serpinh1 | serpin peptidase inhibitor, clade H (heat shock protein 47), member 1, (collagen binding prote | 17493658 | 0.581 | 8.47E-04 | other | yes | yes | |
| Sgtb | small glutamine-rich tetratricopeptide repeat (TPR)-containing, beta | 17289553 | 0.534 | 1.51E-03 | other | | | |

**Correlation between datasets**

| Symbol | Entrez Gene Name | Affymetrix ID | Fold change | p value | Type | Fibrosis association | Inflammation association | Metabolism association |
|---|---|---|---|---|---|---|---|---|
| Hmga1 | high mobility group AT-hook 1 | 17335129 | 0.592 | 1.23E-02 | transcription regulator | | | |
| Rab20 | RAB20, member RAS oncogene family | 17507435 | -0.508 | 4.75E-03 | enzyme | yes | | |
| Rhbdl2 | rhomboid, veinlet-like 2 (Drosophila) | 17418223 | 0.635 | 7.04E-03 | peptidase | | | |
